# Deep spatial-omics to aid personalization of precision medicine in metastatic recurrent Head & Neck Cancers

**DOI:** 10.1101/2023.02.10.527955

**Authors:** Andrew Causer, Xiao Tan, Xuehan Lu, Philip Moseley, Min Teoh, Margaret McGrath, Taehyun Kim, Peter Simpson, Christopher Perry, Ian Frazer, Benedict Panizza, Rahul Ladwa, Quan Nguyen, Jazmina L Gonzalez-Cruz

## Abstract

Immune checkpoint inhibitor (ICI) modality has had a limited success (<20%) in treating metastatic recurrent Head & Neck Oropharyngeal Squamous cell carcinomas (OPSCCs). To improve response rates to ICIs, tailored approaches capable to capture the tumor complexity and dynamics of each patient’s disease are needed. Here, we performed advanced analyses of spatial proteogenomic technologies to demonstrate that: (i) compared to standard histopathology, spatial transcriptomics better-identified tumor cells and could specifically classify them into two different metabolic states with therapeutic implications; (ii) our new method (Spatial Proteomics-informed cell deconvolution method or *SPiD*) improved profiling of local immune cell types relevant to disease progression, (iii) identified clinically relevant alternative treatments and a rational explanation for checkpoint inhibitor therapy failure through comparative analysis of pre- and post-failure tumor data and, (iv) discovered ligand-receptor interactions as potential lead targets for personalized drug treatments. Our work establishes a clear path for incorporating spatial-omics in clinical settings to facilitate treatment personalization.

## Introduction

Immunotherapy and more specifically check-point inhibitors (ICIs) have revolutionized the management of solid tumors but despite the great advancements, ICIs have been limited by a lack of ability to confidently identify patients who are likely or unlikely to positively respond to treatment ^1^ A prime example is mucosal Head and Neck Squamous Cell Carcinomas (HNSCCs) where multiple studies have failed to show a benefit of ICIs over standard chemotherapy ^2–4^. This failure is partially explained by the high inter- and intra-tumor heterogeneity of mucosal HNSCCs with different tumor microenvironments (TME) influenced by the location of primary site and etiology (i.e., HPV^+^ *vs* HPV^-^) ^5^. Although there are over 4,000 active clinical trials testing the efficacy of PD-1/PD-L1 ICIs +/-additional agents and 300 trials for new targets, only a few predictors for their benefit in treating metastatic recurrent HNSCCs are available (i.e., high PDL-1 levels in the tumor) ^6,7^. Therefore, for ICIs to reach their full therapeutic potential the oncology field needs to develop comprehensive analytical pipelines to categorize patients based on their tumor’s likelihood to respond to a specific treatment. The advent of new technologies has provided the knowledge that not only expression but also distribution of targets and interactions between malignant and non-malignant cells (immune system and stroma) can predict response to treatment, tumor development and progression ^8–10^. Recent development in spatial-based high-throughput technologies, such as spatial transcriptomics (ST) and spatial proteomics (SP), have the ability to assess information of cell subpopulations whilst maintaining the spatial architecture of the tissue, thus providing an unprecedented level of knowledge about complex biological systems including tumor development and response to treatment ^11^. Here, as illustrated by a case example, we show how spatial proteogenomic data can make precision medicine a reality by rapidly resolving patient’s disease heterogeneity and generating quantitative data that informs about druggable targets, which alone or in combination have the highest likelihood of delivering a desired therapeutic response.

## Results

### Spatial transcriptomic mapping of tumor and healthy tissue comprehensively distinguish cell distribution and composition at a level not achievable by traditional methods

While methods like MRI and PET-CT scanning can capture general changes in size and locations of the tumor, deeper analysis of the cancer cells, expression of drug targets, and tumor microenvironment is required for a more accurate view of disease status (**Figure 1A-B**). Due to its ability to produce whole-transcriptome resolution (>22,000 transcripts per tissue section) while maintaining spatial information and tissue morphology (a histopathological image accompanied by pathological annotation), spatial transcriptomic 10x Chromium Visium (ST) was chosen to analyze MAR21 OPSCC tumor and a healthy soft palate control sample. Following data quality control and integration (*Supplementary appendix*), unbiased clustering based on gene expression profile similarity (DEG) identified 11 distinct clusters that closely recapitulated the tissue architecture (**Figure 1D**). We annotated each spot cluster using the manually curated JENSEN tissue-gene association reference database (V2.0) (**Figure 1E**). Transcriptional-based annotations matched those independently supplied by the pathologists (**Figure 1F**), whereby cluster 4 (CL4) and 5 (CL5) overlayed the tumor sites (**Figure 1D, F**). In addition to the main cancer clusters, other cell/tissue types were annotated, providing a comprehensive view of the entire tissue section, including epithelium (CL3), muscle (CL7), blood vessels (CL10) and pharynx (CL2) (**Figure 1F**). Of note, carcinoma clusters were annotated as the cervical epithet due to the HPV^+^ OPSCC gene signature commonalities with cervical cancer and overrepresentation of the former disease in the JENSEN database ^12^. Thus, cancer clusters (annotated as “Cervical” carcinomas) orientated in a nest-like structure (differentiated OPSCC), with CL4 being the edge and CL5 the core of the tumor. In contrast to CL5, exclusively found within the tumor biopsy (**Figure S1**), 30% of CL4 was located within the healthy tissue (**Figure 1D, E; Figure S1**). In depth analysis of CL4 facilitated by the heterogeneity between spots in the ST data, allowed us to re-annotate spots in CL4 into three categories, whereby the spots present in both the healthy tissue and the tumor were confirmed as epithelium, whereas spots that were annotated as carcinoma were only present in the tumor (**Figure 1G**). These results highlight the heterogeneous nature of the tumor biopsy, and the capability of ST to successfully distinguish between tumor regions and healthy epithelial tissue based on transcriptional profiles. Most importantly, ST identified different tumor niches visually undistinguishable, but with clear metabolic signatures enriched in specific biological functions such as CD4^+^ T-cell activation (CL4) and angiogenesis (CL5). Such in depth source of information not achievable by standard pathological annotation is crucial to accurately assess the nature of the disease and the potential effects of drugs on various cell types across the tissue.

**Figure 1.**
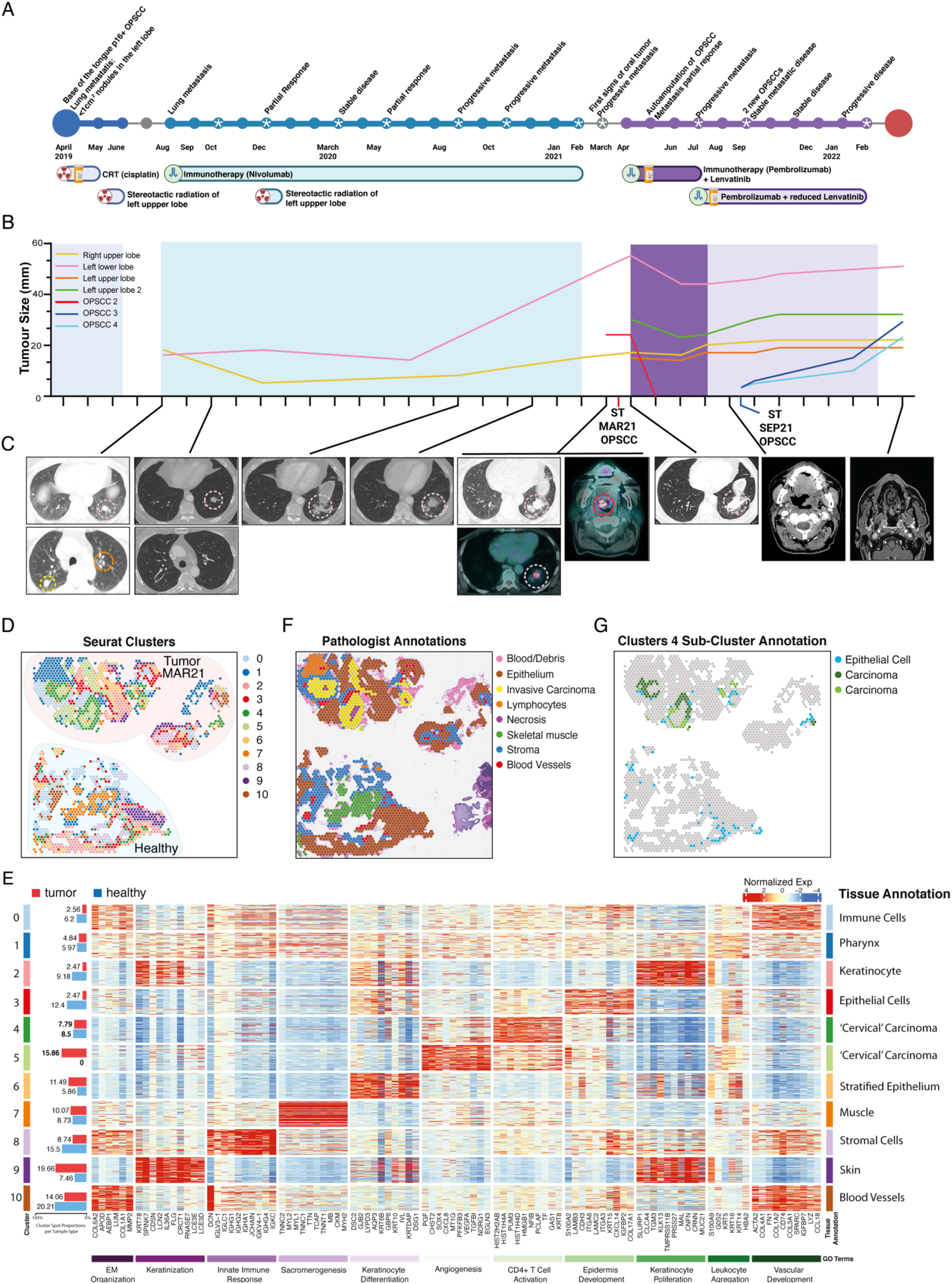
Case history and preliminary ST analysis of metastatic recurrent HPV^+^ OPSCCs. **A.** Timeline indicating disease progression and treatment history. **B.** Relative size and development of OPSCCs and lung metastases over time. Red line indicates the MAR21 soft palate OPSCC sample biopsied for Visium ST and Phenocycler SP analysis. Dark blue line represents recurrent SEP21 OPSCC analyzed using Visium ST. **C.** CT scans of lung metastases and OPSCCs over time. Circled regions highlight tumor tissue. **D.** Spatial representation of unsupervised ST-generated clustering results. Colored spots represent different populations of spots that share similar transcriptional profiles. **E.** Normalized expression values of the top 10 distinguishing markers for each cluster were displayed in a heatmap. JENSEN TISSUEs annotations and relative enriched Gene Ontology (GO) terms associated with differentially expressed genes (DEGs) for each cluster are also represented. Bar graph (left) represents the proportion of each cluster found within MAR21 tumor (red) and healthy (blue) samples. **F.** Pathologist annotations of MAR21 and paired healthy sample, defining general tissue structures including stroma (blue), skeletal muscle (green) and invasive carcinoma (yellow). **G.** Sub-clustering of cluster 4 identified distinct carcinoma sub-clusters which were localized only within the tumor sample (green and dark green). Blue spots represent the third sub-cluster (cluster 4.Epi) which was annotated as epithelial tissue based on DEGs (using JENSEN TISSUEs database).

### Unbiased, transcriptome-wide analysis of spatial gene expression identifies two distinct tumor microenvironments

Collectively, over 5,000 genes out of >22,000 genes were significantly over-expressed in the CL4 and CL5 with signatures enriched for gene ontology terms associated with mRNA processing and transport, DNA regulation and repair, and cell cycle regulation (**Figure 2A**). Sustained proliferation is a hallmark of cancer, and mitoses are used for diagnosis and to grade these malignancies ^13^. Clinically, targeting vital cancer pathways activated to avoid mitotic catastrophe have proven to be of great therapeutic value ^14^. Therefore, we investigated the spot’s proliferative status based on the expression of cell-cycle-related genes (**Figure 2B**). Previously defined clusters showed different proportions of spots in each G1, S and G2M phase. “Skin-related” (sharing epithelial origin) clusters such as CL2, 6 and 9 showed the least proliferative profiles (**Figure 2C**). Remarkably, of all 11 clusters, the CL4 was the only one that contained 100% spots in the proliferation phase (S with 45.3% and G2M at 54.7%), indicating active cell division and expansion of this cancer cluster. As the cluster mapping and histopathological features suggested that CL4 and CL5 form two layers, with core (CL5) and peripheral (CL4), we then sought to analyze the genes that were differentially defining tumor CL4 and CL5 (**Figure 2D**). CL4’s differentially expressed genes (DEGs) confirmed the proliferative nature of the cluster with upregulation of *HIST1H* family genes, which participate in nucleosome assembly and chromatin organization, and *GABRP* which promotes cell proliferation in oral SSC models (**Figure S2**). Conversely, CL5’s DEGs were enriched with genes involved in innate immune response, inflammatory processes, cell migration and angiogenesis, such as *CXCL8*, which attracts neutrophils, basophils, and T-cells or *S100A7*, involved in activation of the innate immune response to viruses (**Figure S2**). The spatial distribution of cell-cycle-related genes allowed us to distinguish 2 distinctive tumor metabolic phenotypes with clinical implications specially when using replicative stress or DNA-damaging agents, as differences in replication can correlate with different response to treatment ^15,16^.

**Figure 2.**
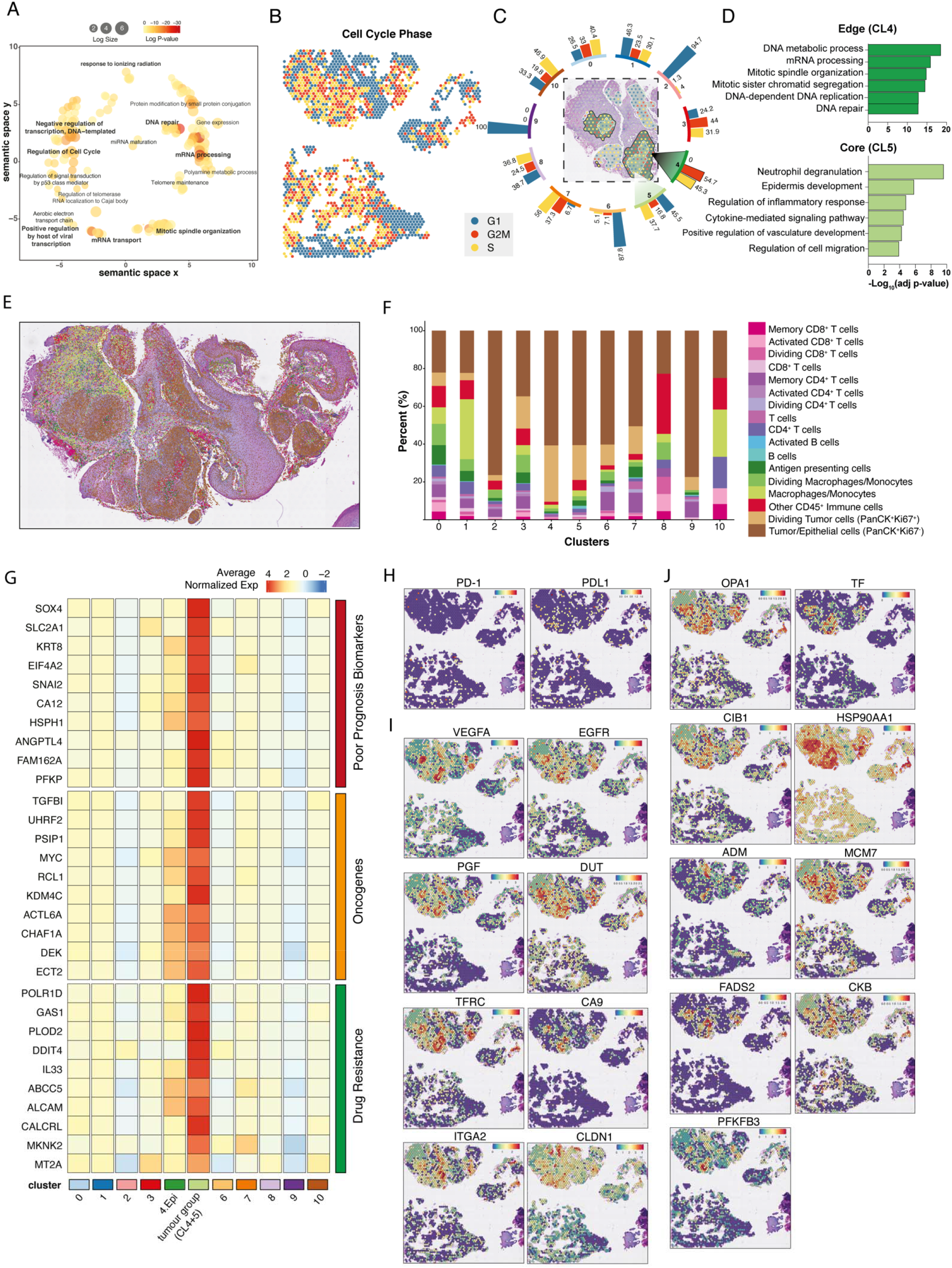
Transcriptional and functional profiles of the tumor microenvironment interface. **A**. Common GO terms enriched in all tumor-annotated spots based on DEGs relative to all other clusters. **B.** Cell cycle states of each spot within MAR21 OPSCC and healthy paired samples. Teal, red and yellow represent spots in G1, G2/M and S phase respectively. **C.** Displays the proportions of spots in each cell cycle phase for each Visium defined cluster. Teal, red and yellow bars represent G1, G2/M and S phase respectively. Distinct tumor regions (cluster 4 and 5) are out-lighted in cluster-matched shades of green (dark green: cluster 4; light green: cluster 5). **D.** Gene ontology terms specifically enriched in each tumor cluster. Significant GO terms were generated based on DEGs between distinct cancer clusters, newly defined Visium clusters 4 (dark green) and 5 (light green). **E.** Single-cell resolution of cell types within the MAR21 sample based on integrated Phenocycler data. Cell phenotype was based on co-expression of marker antibodies. **F.** Stacked bar graphs highlight the percentage of different Phenocycler cell subtypes observed within each Visium cluster. Colors are paired to indicate similar cell types (e.g., pink and purple are T cell subtypes, blue shades are B-cell subtypes). **G.** Functional classification of DEGs (p<0.001) of the combined tumor clusters relative to all other clusters and at least 2-fold overexpressed (CL4 and CL5). Relative expression of poor prognosis markers (red), oncogenes (orange) and drug resistance genes (green) across each cluster. The top 9 genes of each category were displayed. Cluster ‘4.Epi’ represents the sub-cluster 4 annotated as epithelial tissue, and ‘tumor group’ represents the carcinoma annotated spots from cluster 4 and 5 combined. Colors gradient represents average normalized expression values across all spots in each cluster, which were z-transformed by genes (rows of the heatmap). **H.** Spatial expression of PD-1/PD-L1 encoding genes across each spot. **I.** Spatial expression of genes targeted by current clinical therapies for various cancer types. **J.** Spatial expression of experimental targets informed by preclinical studies. Both clinical and preclinical druggable targets were identified based on genes significantly overexpressed (p-value < 0.001 and > 2-fold change) within the grouped tumor clusters.

### Integration of spatial proteomics data enabled the mapping of 14 immune subtypes to each spatial transcriptomics spot, further confirming distinct cancer microenvironments

As infiltration and localization of immune cells in the TME are biomarkers of disease progression and treatment outcome, we sought to identify the cell composition of each cluster in the tumor biopsy at single-cell resolution. Although Visium data produces high-resolution transcriptional data, each spot is a mixture of on average 1-9 cells. Typically, spot deconvolution is needed to identify the cell proportion in each spot. Since no ideal deconvolution method is currently available, we compared the performance of 4 established deconvolution methods (STdeconvolve, CARD, Seurat label transfer and RCTD)^17–20^. All these methods solely use transcriptomic data that comes from reference datasets or directly inferred by the tissue transcriptome. Importantly, to achieve single-cell resolution and subclassification of immune cells, we devised a new method to integrate Phenocycler spatial proteomics (SP) data with ST data. By mapping consecutive tissue sections, we were able to infer the signal of multiple lineage antibodies to map cell types to Visium spots (**Figure 2E, S3**). Overall, our approach identified 14 different immune cell subsets, including T-cells, B cells, macrophages and antigen presenting cells, and allocated the spatial location of specific immune cell types, such as CD4^+^ T-cells and activated B cells, that were not identified using the other 4 indirect cell deconvolution methods. All deconvolution methods overestimated the proportion of cancer cells and were insensitive in detecting immune infiltration (**Figure S3**). Therefore, our Phenocycler SP-informed cell deconvolution method (*SPiD*) outperformed the four most used spot resolution methods.

### High resolution cellular composition of the tumor defined by integrating SPiD suggests tumor functional organization

*SPiD* deconvolution confirmed our cell-cycle prediction at the protein level by detecting high expression of Ki67 (S-G2/M phase surrogate) in the tumor nests (intra-tissue PanCK^high^) (**Figure S4**). Once more, CL4 recorded the highest proportion of Ki67^+^ PanCK^+^ cells (29% of CL4) (**Figure 2F**, **Figure S5**) forming the walls of the tumor as previously seen in the Visium ST data (**Figure 2C**). Conversely, CL5, composed by the inner core of the tumor, displayed lower proportions of dividing tumor cells (18.4%) and higher levels of immune cells (21%) compared to CL4 (9.5% immune cells). Tumor CL5 displayed the largest proportions of activated and non-activated B cells across all clusters. This inner tumor core also contained high levels of macrophage/monocytes and CD4^+^ and CD8^+^ T-cells. Collectively, our spatial proteo-transcriptomics data confirmed the presence of two functionally distinct tumor regions representing the proliferating leading edge and immune infiltrated inner core of the patient’s OPSCC which phenotype and cell-type complexity could not be resolved by histopathological assessment (**Figure 1E**).

### The spatially defined tumor microenvironments informed the assessment of predictive biomarkers and druggable targets

Traditional methods such as bulk RNA sequencing suffer from dilution of cancer specific signals within the pool of non-malignant tissue. We hypothesized that our ST data would over-come this by allowing us to focus on the most important part of the tumor, namely the proliferative cancer tissue depicted by only 2 clusters out of the 11 that composed the biopsy. This spatially-focused analysis strategy maximizes resolution while minimizing signal dilution as seen in bulk data analysis. By doing so, we identified 158 significantly up-regulated (p < 0.001) tumor genes which displayed at least a 2-fold increase compared to all other clusters (**Figure S6**). The majority of top genes were categorized as poor prognosis biomarkers (38%), oncogenes (13%) and drug resistance genes (9%) in contrast to only a 11% and 5% considered tumor suppressor and positive prognosis biomarkers, respectively (**Figure 2G, Table S3**). Notably, the function of 6% of the >2-fold increased tumor genes remain unknown, and hence may potentially be new markers. These results correlate with the observed recurrent and aggressive behavior of the patient’s tumor and highlights the potential of focused transcriptional profiles to predict disease progression.

The patient’s treatment history pinpoints two specific proteins targeted by immunotherapy and chemotherapy: PD-1 (Nivolumab, Pembrolizumab) and VEGFR (Lenvatinib). Interestingly, we found that both PD-1 and its ligand PD-L1 encoding genes displayed low expression across the whole tissue (**Figure 2H**). Indeed, these two targets were not expressed in the two cancer regions, potentially explaining why ICIs failed. Conversely, amongst the 158-tumor overexpressed genes there were 8 targets commercially available or in clinical trials, including *EGFR*/cetuximab-prochlorperazine (**Figure 2I**) ^8^ and 9 experimental targets supported by preclinical data (**Figure 2J, Table S4**) ^21^. Consequently, the use of ST has the potential to prevent the use of treatments that are unlikely to be effective while simultaneously identifying novel therapeutic targets.

### Tumor clusters and targets were shared by pre- and post-treatment tumors and revealed potential causes of treatment failure/response

In the recurrent settings, it is imperative to investigate whether information obtained from a tumor can predict the phenotype of tumors to come. Thus, we spatially sequenced the SEP21 OPSCC tumor (failed combinational therapy) and compared it to the previously sequenced MAR21 tumor (failed monotherapy) (**Figure 3A**). Annotations based on transcriptional profiles were consistent with pathologists’ evaluation, showing that ST was capable of successfully identifying tumor cells (**Figure 3B**). Furthermore, transcriptomic annotation aided the pathology team in resolving a conflictive area where low stroma abundance made it difficult to assess the invasive cancer (tumor), highlighting the power of using unbiased molecular profiles to characterize tissue regions (**Figure S7**). To compare cancer cells in both biopsies, we performed the same cell-cycle analysis (**Figure 3C**) and unbiased gene markers identification for clusters (**Figure 3D**) as for MAR21 (Figure 1D). The consistent results mapped the same carcinoma hubs in both biopsies, where MAR21 CL4 and CL5 correspond to SEP21 CL4 and CL7 (**Figure 3D, S8**). Importantly, most of the identified MAR21 druggable and experimental targets were also over-expressed by SEP21 tumor clusters (**Figure 3E-G**, **S8**, **Table S3**, **S4**). These druggable targets were identified in separate tissues and at different time points, suggesting the reproducibility of detecting potential targets, which colocalize to cancer regions and maintain a high expression level throughout time and space.

**Figure 3.**
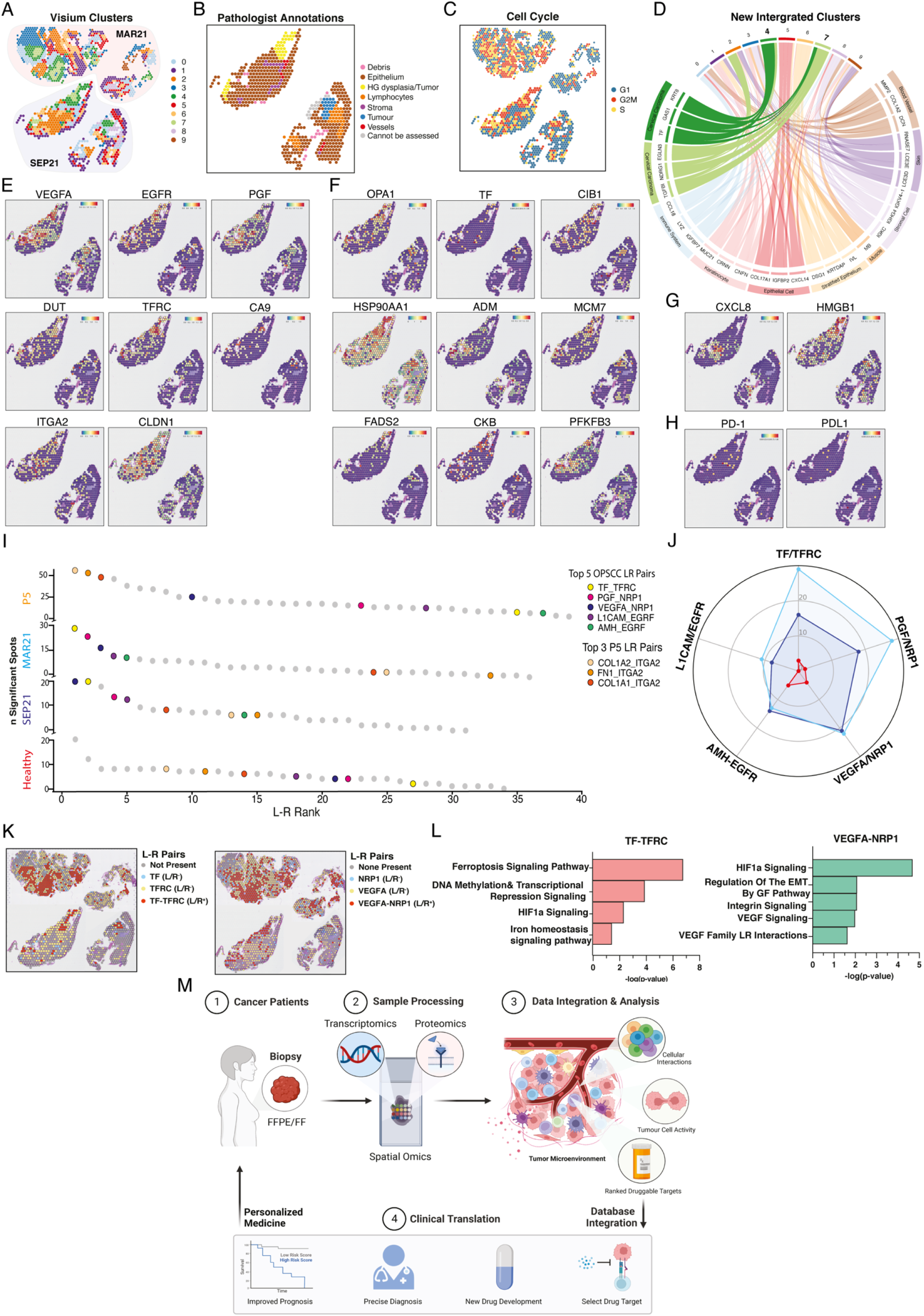
Transcriptional comparison between MAR21 and recurrent SEP21 and Ligand-receptor interaction analysis for therapeutic target selection. **A.** Spatial representation of unsupervised clusters identified between integrated tumor samples. MAR21 highlighted in red shades and recurrent tumor SEP21 highlighted in blue shades. **B.** Annotated cell cycle phase of each spot based on relative expression of specific cell phase genes. **C.** Pathologist annotations of SEP21, defining general tissue structures including epithelium (brown), dysplastic tissue (light blue) and tumor (dark blue). **D.** Chord diagram displays genetic correlation between new clusters corresponding to integration of the MAR21 and SEP21. Comparisons were based on gene expression levels of the top 3 upregulated genes expressed by each original MAR21 cluster (only one muscle gene was found in the new cluster) within each new cluster. Connecting ribbons define the correlation between original and new clusters which significantly over-express each gene. **E.** Spatial expression of genes across each spot targeted by current clinical therapies. **F.** Spatial expression of genes targeted by select experimental drug therapies. **G.** Spatial expression of genes targeted by select experimental drug therapies only seen in SEP21 samples. **H.** Spatial representation of gene expression for Nivolumab and Pembrolizumab targeted *PD-1/PD-L1* pathway. **I.** Ranking of top 35 Ligand/Receptor (L/R) pairs targeted by clinical and experimental therapies expressed by the healthy (red), MAR21 (red) recurrent SEP21 (blue) and additional patient (orange) samples. Rank was based on the number of significant spots expressing each L/R pair across each sample. Colors indicate the location of each specific LR pair. Orange spots represent the top 3 L/R pairs specific to the additional sample. **J.** Spider-plot represents the top 5 most active L/R pairs found across tumor tissue based on the number of spots co-expressing both molecules. **K.** Spatial localization of ligand and receptor expression across tumor tissues (top: MAR21, bottom: SEP21). Red spots indicate co-expression of both ligand and receptor within the same spot. **L.** Significantly active IPA pathways associated with expression of the top two ranked L-R pairs (TF/TFRC and VEGFA/NRP1). Bars indicate negative-log p-value of each pathway based on the proportion of genes within each canonical pathway that were also significantly up- or down-regulated within spots co-expressing both L-R pairs. **M.** Infographic representing the proposed protocol to incorporate spatial multiomics within a clinical setting.

The initial lack of response to anti-PD-1 treatment and subsequent resurgence of a new locoregional tumor suggests evasion by a drug resistance mechanism. The deep analysis of spatial proteo-transcriptomics of the MAR21 tumor showed no expression of PD-1 in the tumor hubs, but rather scattered expression of this target across the tissue section in non-cancer cells, potentially being the cause for the lack of response. In contrast, the combinatorial therapies targeting PD-1 and VEGFR at the same time showed initial responses with tumor self-amputation, followed by tumor resurgence when VEGFR was reduced to aid with patient’s coagulation/healing. Analysis of the SEP21 sample showed low PD-1/PDL-1 and high *VEGFA* levels at the tumor locations. Overall, our findings strongly suggest that the therapeutic response to MAR21 OPSCC was not driven by Pembrolizumab but lenvatinib, as its reduction caused the growth of recurrent VEGFA^high^ tumors previously suppressed by the chemotherapy.

### Novel method to prioritize targets based on Ligand/Receptor interactions

Treatment personalization is considered the future of many medical disciplines including oncology. As many targeted anti-cancer drugs act by blocking ligand/receptor interactions (L/R), we explored the possibility of tailored treatment directed against patient’s tumor core upregulated genes by ranking the relative functionality of these targets based on the colocalization and activity of L/R pairings in the tumor microenvironment (**Figure 3I**). Based on the expression levels of co-localized L/Rs in each sample (healthy, MAR21 and SEP21), we implemented *stlearn* methods ^22^ to rank the 50 L/R pairs that are known to be potential genetic targets of previously identified clinical and preclinical drugs (**Figure 2I-J, Table S3-4**). This classification allowed us to select the top-5 most active L/Rs in the tumor *vs* the healthy tissue, being *VEGFA/NRP1* and *TF/TFRC* highly active in both the original and recurrent OPSCCs (**Figure 15J**). Importantly, spatial analysis identified the coexpression of these L/R pairs primarily within the malignant regions of both tumor samples, which would specifically focalize the treatment to the malignant tumor clusters (**Figure 3K**). The selected targets were patient-specific, as the analysis of another individual’s HPV^+^ OPSCC, showed that although some interactions, such as *VEGFA/NRP1* were present, the most active L/Rs in the tumor were *ITGA2*-related pathways (**Figure 3I, S9**). The analysis of the additional patient sample showed that this patient had a high expression of PD-1 and PD-L1 in the cancer core region, a contrasting pattern compared to the case reported here, suggesting that this patient might be responsive to PD-1/PD-L1 drug. Thus, spatial gene expression profiling can detect specific expression patterns of drug targets in each patient sample. Lastly, we confirmed that both *VEGFA/NRP1* and *TF/TFRC* interactions were biologically functional as DEGs from L/R-positive *vs* the L/R-negative spots were significantly enriched in canonical pathways associated with the downstream activation of these L/R pairs (Ingenuity Pathway Analysis (IPA), **Figure 3L**). Overall, we identified that L/R co-expression and downstream pathway analysis can be used to prioritize clinical targets allowing clinicians to combine multiple strategies that will interfere with known tumor-activated pathways in a personalized manner.

## Discussion

Although the value of the global personalized medicine market is expected to reach USD 922.72 billion by 2030, a 6^th^ of which will account for Oncology Precision Medicine alone ^23^, very few biomarkers are capable of predicting cancer response to treatment with reasonable certainty. In fact, tumor heterogeneity and constant disease evolution indicate that patients will only benefit long-term from tailored medicine if they are based on analytical pipelines that are as holistic and dynamic as the disease itself is. Studies using spatial-omics techniques to understand cancer disease development and progression are increasingly more common in the field of oncology research ^24^. Here, we sought to explore the use of spatial multi-omics in a clinical context, to determine its appropriateness as a potential medical tool for recapitulating patient disease progression and aiding in the drug selection and combination processes.

Based on spatially defined, differentially expressed genes, we resolved intra-tumor heterogeneity and confidently annotated diverse tissue types including malignant and healthy, stroma and infiltrating immune cell populations. Two clusters (CL4 and 5), which overlayed the cancerous and necrotic regions defined by pathologist annotations, were highlighted by multiple lines of unbiased analyses as carcinoma tumor communities showing differential metabolic signatures: CL4, highly proliferative leading-edge *vs* CL5, core enriched in immune infiltrates interacting with tumor cells (**Figure 1**). Detection of two cancer clusters, with distinct molecular, but not morphological phenotypes, was an important result of our data-driven approach. Although, Ki67 protein detection confirmed the proliferative nature of CL4, expression of cell-cycle-phase associated genes predicted a higher degree of cells in S-G2M phases (**Figure 2C**). This suggests that ST method was more sensitive at detecting dividing cells by assessing hundreds of cell-cycle-related genes in contrasts to a sole marker as immunohistochemistry methods usually use ^13^. Therefore, this method could aid the assessment of the tissue mitotic activity in an unbiased manner, pinpointing regions with heterogeneous metabolic profiles that could impact prognosis and response to treatment targeting replicative stress. Importantly, the cytotoxicity of certain chemotherapies such as Cisplatin and Paclitaxel, depend on the tumor proliferative state ^15,16^. For instance, repeated dosages of Paclitaxel were more efficient when given in cell culture to cells preparing for G2M phase ^16^. ST data of core biopsies taken at different time-points during the treatment could be an excellent tool to test whether timing optimization of sequential chemotherapeutics improves their cytotoxic capacity in a clinical setting.

To be noted, all analyses performed were conducted prior to receiving tissue annotations, emphasizing the true unbiased and discovery strengths of our approach and findings. For the second tumor (SEP21), the transcriptomic signature also informed pathologists of tissue areas with suspicious features, which facilitated the resolution of conflict zones resulting in a more assertive annotation of the dysplastic regions. We have proven that unbiased annotation based on ST data recapitulates clinical annotations and can assist pathologists especially when the quality/size of the sample is not optimal for visual macroscopic characterization. Although, more data needs to be collected and included in reference databases to enhance the accuracy of the annotations (i.e., HPV^+^-cervical cancer *vs* HPV^+^-OPSCC), overall, the ability to precisely define these cancer regions and assess specific markers differentially expressed by them has a great potential for drug selection.

The interactions between myeloid cells and lymphocytes with tumor cells play an important role in the functioning TME and can determine treatment outcome ^25,26^. Such interactions ideally require single-cell resolution data and detailed identification of immune cell types. However, the current Visium ST technology suffers from relatively low single-cell resolution as ST ‘spots’ encompass average gene expression across several cells (1-9 cells). To address this issue, we developed a novel spatial multi-omics deconvolution approach (*SPiD*) by integrating Visium ST and Phenocylcer SP data which outperformed the currently 4 most used methods. Importantly, our method has the additional advantage of considering both protein and transcriptional data when studying tumor and TME interactions, which increases the robustness of cell identification, especially of immune cell populations, by overcoming the problem of the non-linear relationship between RNA expression and protein levels ^27–30^. Using our protein/RNA-based method we identified 14 different immune subtypes including activated CD4^+^ and CD8^+^ T cells, macrophages and B cells present within the inner tumor, which would otherwise be impossible by sole visual evaluation of the sample. Our method enables the association of the cell type to their corresponding transcriptional profile. This information can correlate cell subsets with disease progression and response to treatment as previously seen for HNSCC, where certain CD4^+^ and CD8^+^ T-cell phenotypes correlated with longer progression-free survival ^31,32^.

The spatial component of our analysis allowed us to focus on the critical tumor (CL4-5), where we observed that most upregulated genes were either oncogenes, poor prognosis biomarkers or drug resistance genes confirming the active and aggressive nature of our patient’s disease ^33^. In the clinic, this information could help stratify patients and tailor surveillance plans based on expected disease behavior. The high-throughput nature of our approach allows answering questions such as, which suitable targets the tumor hubs are expressing. Although PD-1/PD-L1 expression was very low, we identified 8 over-expressed druggable targets (*i.e., EGFR, TF, VEGF*) and 9 preclinical targets in CL4-5. Although, more work is needed to prove the robustness between the presence/distribution of a target and therapy response, being able to interrogate the patient’s tissue whether a target is present or not will save time, resources and psychological burden to patients and their families. In fact, these drugs and downstream analysis results were patient-specific, reinforcing the real need for tailor approaches. Nonetheless, targets were mostly shared by the recurrent tumors of the same area, indicating that in this case the information gained from the first patient’s OPSCC biopsy still applied to his potentially preventable subsequent malignancy. However, considering the time and health constraints of recurrent non-responsive cancer patients, a list of targets might not guarantee patient’s long-term clinical response, thus a way to rank the target candidates is equally vital. We reasoned that the likelihood of a drug having an impact on tumor growth and progression would be linked with its capacity to interfere with vital and active pathways in the malignancy. Therefore, we conceived a novel way to rank each drug’s potential success, based on the co-expression of each target ligand-receptor pair (L/R), assuming that co-expression would lead to interaction hence pathway activation. After our analysis, our patient’s list of 17 clinical/preclinical targets was reduced to 3 top druggable pathways active in the patient’s tumor clusters: *TFRC, NRP1* and *EGFR*. Of note, *NRP1* is a SARS-CoV2 receptor, for which dozens of new drugs have recently been developed ^34,35^, and hence may justify further clinical applicability in cancer.

Overall, our work clearly demonstrates the power of Spatial proteogenomic data to resolve tumor intra- and inter-heterogeneity and enables the real possibility for oncologists to personalize cancer management. We envision that spatial RNA/protein analysis can be adopted in the clinical settings, as whole genome sequencing is now a routine test requested by clinicians. Importantly, the cost of these technologies and high technical requirements can already be drastically reduced by the implementation of artificial intelligence (AI) models capable to predict *in situ* gene expression inferred from fast/low-cost H&E images using curated disease Spatial-omics training data sets ^36,37^ Thus, after an initial investment dedicated to create standardized disease-specific AI training material, the spatial data of each patient’s tumor biopsy can be obtained (experimentally or AI-inferred) and contrasted against spatial databases of the disease to help with different steps along each patient’s journey: (i) aid in the annotation of the tumor, (ii) stratify patients based on disease risk progression to personalize surveillance plans; and (iii) to generate a list based on a patient’s own disease features which informs oncologists of targets with quantifiable likelihood to have an impact on the disease. This information can be used to implement combinatorial therapeutic programs to prevent drug resistance minimizing off-target effects (**Figure 3M**).

## Methods

### Clinical history of the patient

A 60-year-old male ex-smoker with a 10 pack-year history presented with *de novo* oligometastatic disease, having been diagnosed with a p16 positive SCC primary tumor of the right tonsil and biopsy-proven left upper lobe lung metastasis (**Figure 1A-B**). He received concurrent chemoradiotherapy comprising weekly cisplatin and 70Gy in 35 fractions of radiation targeting the primary tumor and bilateral level II and III neck nodes. Additionally, he received stereotactic radiation to the left lung nodule of 50Gy in 5 fractions, completed in June 2019.

The patient underwent disease re-assessment three months later transfer of care, with PET/CT demonstrating multiple new bilateral pulmonary metastases and no locoregional disease. He commenced treatment of 480mg nivolumab every 4 weeks and a CT scan after three cycles revealed resolution of two nodules and significant reduction in a third but minor growth of a left lower lobe nodule. This single nodule was then treated with stereotactic radiation of 48Gy in four fractions and nivolumab was continued. Regular imaging confirmed stable intrathoracic disease for a further 13 months but a PET/CT scan in February 2021 demonstrated local recurrence of disease with a 22mm lesion of the left soft palate (biopsy MAR21) and progression of metastatic lesions in the lungs bilaterally.

The patient was then enrolled in a clinical trial (LEAP-009) and randomized to treatment of pembrolizumab 200mg every 3 weeks and 20mg lenvatinib orally daily. After demonstrating an early partial response radiologically, including autoamputation of the oropharyngeal recurrence with absence of measurable disease and a reduction in lung metastases, a treatment delay was required due to oropharyngeal pain and severe non-healing ulceration of the oropharynx. This subsequently settled and the patient recommenced pembrolizumab and dose-reduced lenvatinib (14mg orally daily) in July 2021. The measurable disease remained stable radiologically until January 2022, although mucosal changes over the tonsillar fossa and soft palate were suspicious clinically from September (biopsy SEP21). Pembrolizumab and Lenvatinib were ceased, and the patient came off trial in early February 2022 when MRI scan confirmed definite disease progression involving the oropharynx.

### Spatial transcriptomics

Five μm sections were taken and multiplexed onto Visium Spatial Gene Expression Slides (10x Genomics). Gene expression quantification per spot was performed following 10X Chromium LIT000128 - Rev B (*Supplementary appendix*).

### Spatial proteomics

A serial tissue section (4μm thick) from the MAR21 FFPE block was taken and analyzed using Phenocycler. Coverslip preparation, antibody conjugation, tissue staining, Phenocycler rendering, and imaging were completed in accordance with Phenocycler manufacturer instructions (Akoya Biosciences User Manual, Revision-C)^38^. Antibodies used for tissue staining and their respective targets are described in **Table S1** (*Supplementary appendix*).

## List of Supplementary Materials

Provided in Supplementary appendix:

**Supplemental Methods**
**Supplemental Table S1** Spatial proteomics: phenocycler antibody panel.
**Supplemental Table S2** Identified target genes of preclinical drugs.
**Supplemental Figure S1** UMAP representation of unbiased Visium clustering of MAR21 and healthy paired samples.
**Supplemental Figure S2** Transcriptional profiles of distinct cancer clusters.
**Supplemental Figure S3** PHENOCYCLER-informed deconvolution outperforms established transcription-based deconvolution methods.
**Supplemental Figure S4** Localization of proliferating tumor cells.
**Supplemental Figure S5** Integrated Spatial-Omic characterization of tumor immune cell microenvironments.
**Supplemental Figure S6** Transcriptional profile of tumor clusters within MAR21 OPSCC.
**Supplemental Figure S7** Initial Pathologist annotation of SEP21 OPSCC sample.
**Supplemental Figure S8** Tumor transcriptional profile recapitulated in recurrent OPSCC.
**Supplemental Figure S9** Top druggable targets differ between additional OPSCC patients.
**Supplemental Table S3** Gene classification based on function reported in the literature in the cancer setting.
**Supplemental Table S4** Identified clinical and preclinical targets.
**References for Appendix**

## Notes

### Competing Interest Statement

The authors have declared no competing interest.

