## Supplemental Appendix for "Deep spatial-omics to aid personalization of precision medicine in metastatic recurrent Head & Neck Cancers"

**Supplementary Appendix Table of Contents**

| **Content** |  | **Page** |
| --- | --- | --- |
| 1. **Supplemental Methods** |  | 1 |
| 2. **Supplemental Table S1** | Spatial proteomics: phenocycler antibody panel. | 4 |
| 3. **Supplemental Table S2** | Identified target genes of preclinical drugs. | 5 |
| 4. **Supplemental Figure S1** | UMAP representation of unbiased Visium clustering of MAR21 and healthy paired samples. | 5 |
| 5. **Supplemental Figure S2** | Transcriptional profiles of distinct cancer clusters. | 6 |
| 6. **Supplemental Figure S3** | PHENOCYCLER-informed deconvolution outperforms established transcription-based deconvolution methods. | 6 |
| 7. **Supplemental Figure S4** | Localization of proliferating tumor cells. | 7 |
| 8. **Supplemental Figure S5** | Integrated Spatial-Omic characterization of tumor immune cell microenvironments. | 8 |
| 9. **Supplemental Figure S6** | Transcriptional profile of tumor clusters within MAR21 OPSCC. | 8 |
| 10. **Supplemental Figure S7** | Initial Pathologist annotation of SEP21 OPSCC sample. | 9 |
| 11. **Supplemental Figure S8** | Tumor transcriptional profile recapitulated in recurrent OPSCC. | 10 |
| 12. **Supplemental Figure S9** | Top druggable targets differ between additional OPSCC patients. | 11 |
| 13. **Supplemental Table S3** | Gene classification based on function reported in the literature in the cancer setting. | 12 |
| 14. **Supplemental Table S4** | Identified clinical and preclinical targets. | 18 |
| 15. **References** |  | 33 |

1. **Supplemental Methods**

*Pathologist Annotations*

High resolution H&E images of samples from two patients were provided to a pathologist for pathological and tissue annotation. Loupe Browser was used by the pathologist to outline and label various morphological features observed within the H&E tissue, which were then converted to a ‘Visium spot’ format. Annotations were blindly performed (the pathologist was not made aware of analysis results) succeeding all other analyses completed in this study.

*Spatial transcriptomics*

Five μm sections were taken and multiplexed onto Visium Spatial Gene Expression Slides (10x Genomics). Following slide incubation (60°C for 2 hours) on a Thermocycle (Bio-Rad C1000 Thermal Cycler), H&E staining, imaging and sequencing library preparation were performed in accordance with the Visium Spatial Gene Expression User Guide (CG000407, CG000408, CG000409 – 10x Genomics). Trimmed FASTQ reads were mapped to the human reference genome (version GRCh38-3.0.0) using *SpaceRanger* (*v1.3.0*) and demultiplexed using 10X Loupe Browser (*v6.1.0*). Sequenced raw read and expression data were processed and overlaid with the H&E image (*Seurat*; *v4.0.5*). Samples were analyzed following a general pipeline that consisted of 1) data filtering and quality control, 2) normalization, 3) batch correction and integration, 4) unsupervised clustering, 5) differential expression analysis and 6) other downstream analyses.

*Spatial proteomics*

A serial tissue section (4μm thick) from the MAR21 FFPE block was taken and analyzed using Phenocycler. Coverslip preparation, antibody conjugation, tissue staining, Phenocycler rendering, and imaging were completed in accordance with Phenocycler manufacturer instructions (Akoya Biosciences User Manual, Revision-C)^54^. Antibodies used for tissue staining and their respective targets are anti-CD20-BX007, anti-PANCK-BX019, anti-CD8𝛼-BX026, anti-Ki67-BX047, anti-CD45RO-BX017, anti-CD3ε-BX045, anti-CD107a-BX006, anti-HLA-DR-BX033, anti-CD4-BX003, anti-CD68-BX015 and anti-CD45-BX021. Probe addition and washing/denaturing steps were performed using the Phenocycler CIM software version 1.30.0.12. Images were collected with the 20x objective (0.8 NA) Zeiss Axio Observer and processed using the Zen Blue v3.2 software.

*Pre-processing of Phenocycler data*

The raw Phenocycler data was processed via QuPath software (version 0.3.2) ^6^. Cells were segmented by using the QuPath function cell detection on Phenocycler DAPI channel with default parameters (minAreaMicrons=2.0, MaxAreaMicrons=500.0, watershedPostProcess=True, smoothBoundaries=True and threshold=2.0). The protein expression intensity was then measured for each segmented cells and exported for further QC. The raw protein expression intensity matrices were filtered by quantile. Cells with total counts lower than 0.05 quantile or higher than 0.95 quantile were discarded to remove the outliers. Filtered matrices were then transformed to logarithmic scale for downstream analysis.

*Phenocycler cell type annotation*

Phenocycler cells were annotated based on the expression of the protein markers. In total, 17 cell types were identified based on the combination of 11 key immune and cancer markers, including tumor (PANCK), tumor infiltrating cells (CD45), Dividing tumor (KI67, PANCK), Dividing macrophage (KI67, CD68), Dividing CD8 T cells (KI67, CD45, CD3e, CD8), Dividing CD4 T cells (KI67, CD45, CD3e, CD4), Antigen-presenting cells (CD45, HLA-DR), T-cell (CD45, CD3e), B-cell (CD45, CD20), Activated B cells (CD45, CD20, HLA-DR), CD8 T cells (CD45, CD3e, CD8), CD4 T cells (CD45, CD3e, CD4), Activated CD8 T cells (CD45, CD3e, CD8, CD107a), Activated CD4 T cells (CD45, CD3e, CD4, CD107a), Memory CD8 T cells (CD45, CD3e, CD8, CD45RO), Memory CD4 T cells (CD45, CD3e, CD4, CD45RO) and Macrophage/monocyte (CD45, CD68). For a given cell *j*, the cell type *Cj* can be defined by the protein marker subset which gives the largest geometric mean value as:

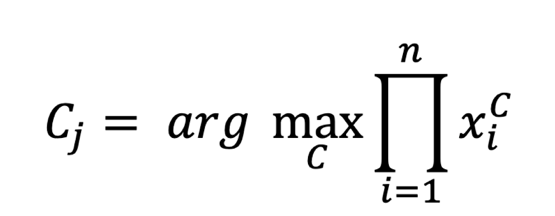

where xiC is the expression value of protein marker *i* that belongs to cell type *C.*

*Image registration analysis of Phenocycler and Visium data*

The Python package SimpleITK was used to perform image registration^6^. Phenocycler images were firstly downscaled to appropriate resolution to match with the resolution of the corresponding Visium histological image. The DAPI channel in the Phenocycler image was cropped and rotated to have the same capture area and orientation with the Visium histological image and was used as the moving image (query image). Visium histological images were converted to grayscale images to transform the pixel data dimension consistent with the Phenocycler DAPI channel image and was used as the fixed image (target/reference image). After centralizing the two images, the rigid affine transformation was applied for shearing, shifting, and scaling the moving image to align with the fixed image in lower resolution as the initial step. Finally, the non-rigid B-spline transformation was applied on affine initialization to refine the local alignment. The mutual information was used as the evaluation matrix to optimize the parameter for both affine and b-spline transformation.

*Integrating Phenocycler protein signal to Visium transcriptional data*

After registering the Phenocycler image to Visium histological image, the optimized transformation matrix was then able to convert the cells in Phenocycler data from the original Phenocycler spatial coordinates (x, y) to newly mapped spatial coordinates (x’, y’) which are identical to Visium spatial coordinates. With this shared coordinating system, cells in Phenocycler data then can be searched and grouped by the spatial radius (d=55um, diameter equivalent to Visium spots size) using the transferred spatial coordinates (x’,y’), which created an additional layer of protein expression profiles on top of existing Visium RNA measurements with compatible resolution (Visium spot level). Those mapped cells were then used to approximately deconvolute the cell type for Visium spots. The proportion *P* for each cell type *C* in a particular Visium spot *s* can be defined as:

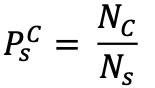

where the N_s_ denotes the total number of cells that fall into Visium spot *s* and of those cells which annotated as cell type C is denoted as *N_C_* .

2. **Supplemental Table S1**

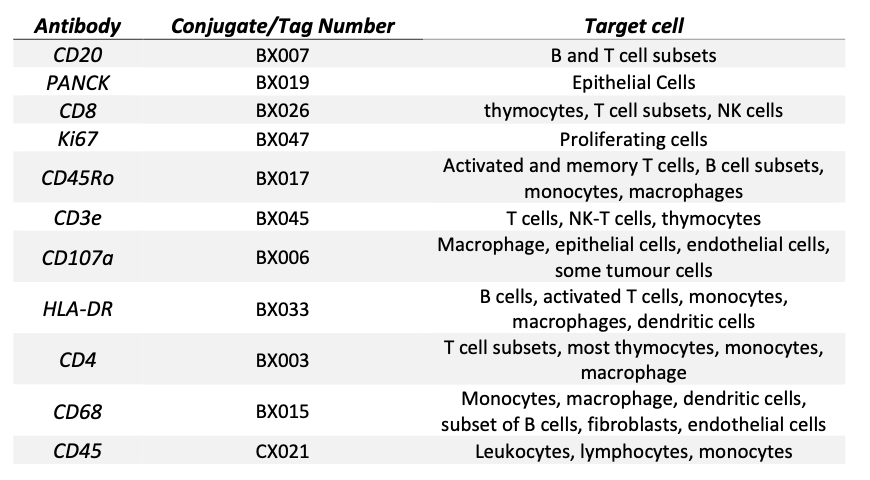

Table S1. **Spatial proteomics:** **phenocycler antibody panel.**

3. **Supplemental Table S2**

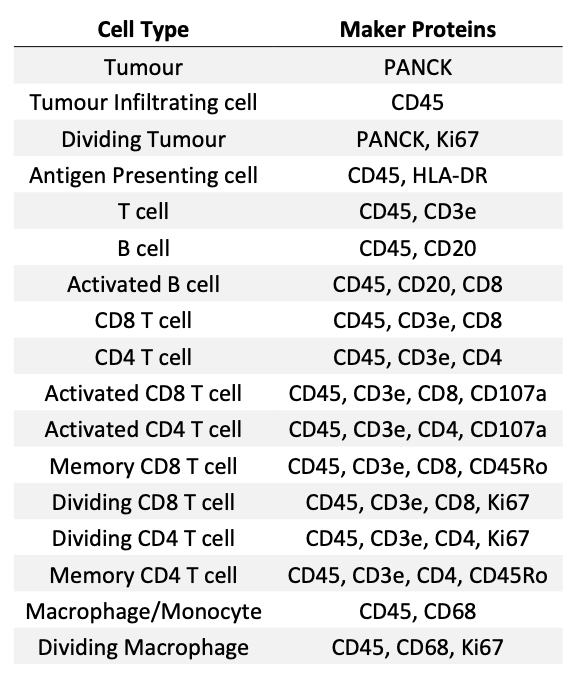

Table S2. **Spatial proteomics: cell type annotation based on surface protein expression.**

4. **Supplemental Figure S1**

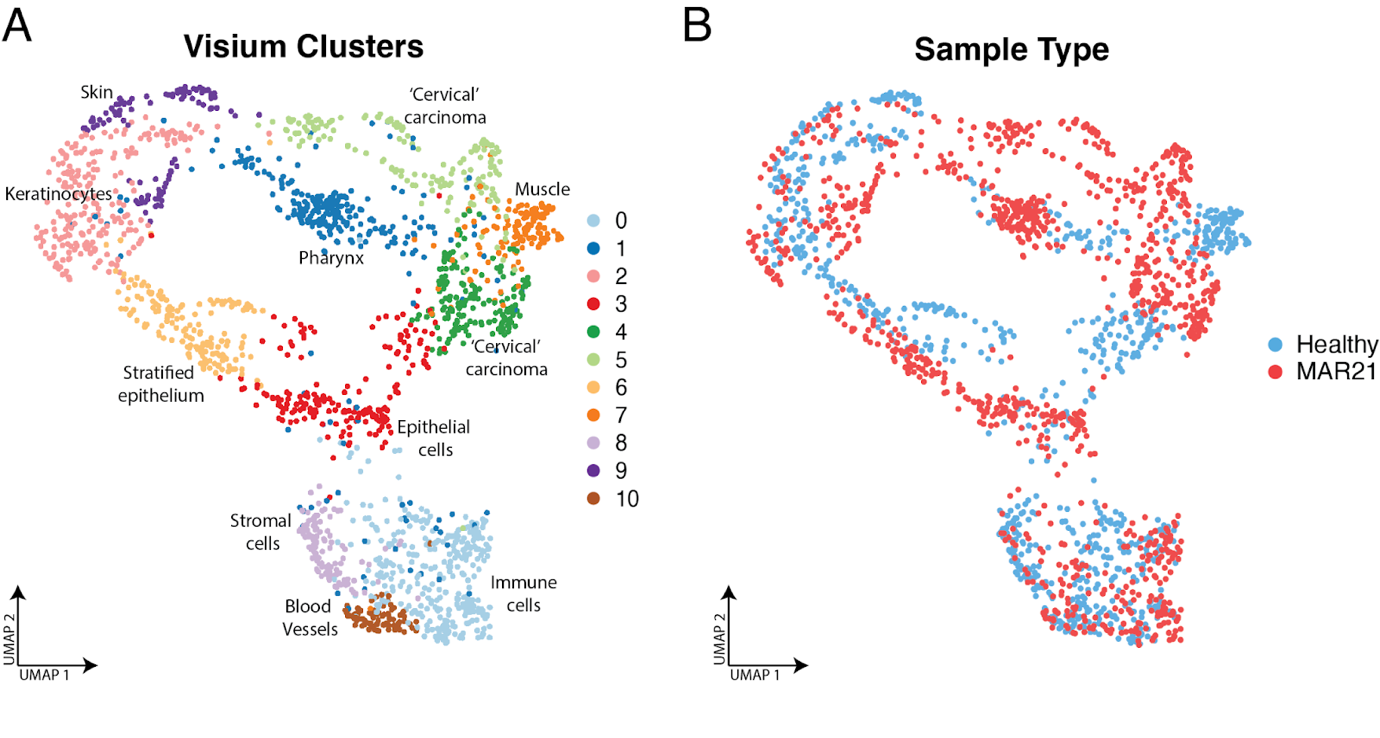

Figure S1. **UMAP representation of unbiased Visium clustering of MAR21 and healthy paired samples**. UMAP of clusters highlights distinction and separation between populations. Cluster annotations were generated by comparing differentially overexpressed genes with a reference database (JENSENs Tissues; EnrichR).  Sample type UMAP displays batch correction between healthy (blue) and MAR21 OPSCC (red) samples.

5. **Supplemental Figure S2**

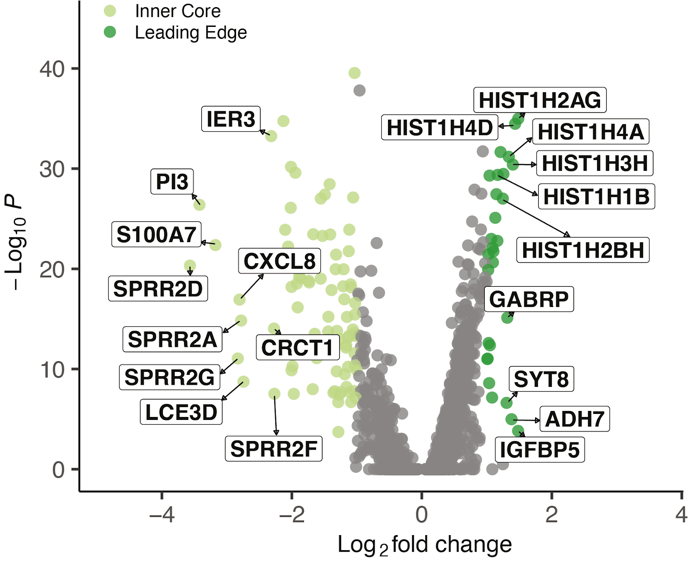

Figure S2. **Transcriptional profiles of distinct cancer clusters.** DEGs between newly defined Visium clusters 4 (dark green) and 5 (light green). Significant DEGs with p-value < 0.001 and fold-change >1 are colored based on cluster.

6. **Supplemental Figure S3**

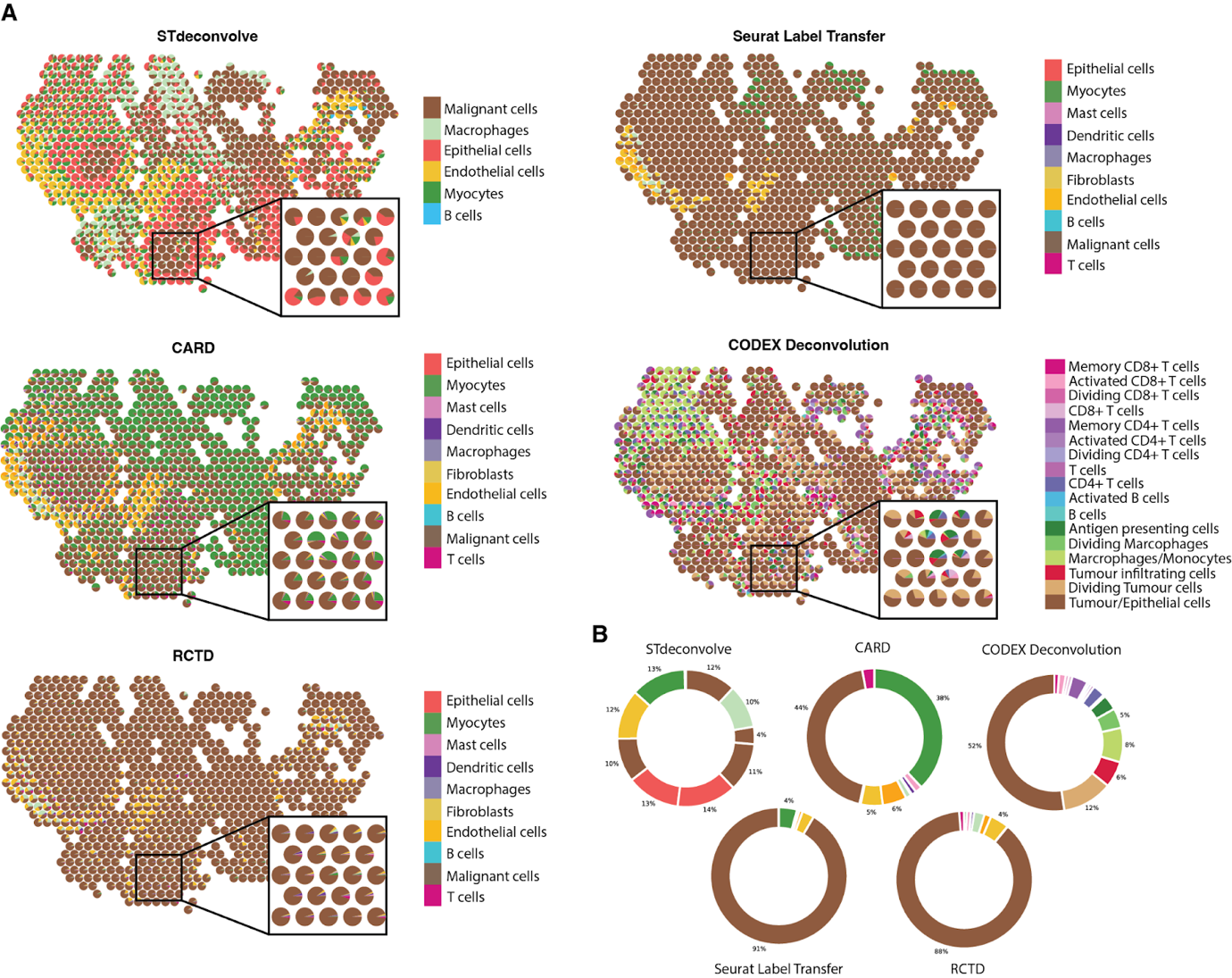

Figure S3. **PHENOCYCLER-informed deconvolution outperforms established transcription-based deconvolution methods. A.** Displays spot deconvolution of tumor sample using each algorithm. Spots are represented as pie graphs denoting the proportions of each cell type deconvoluted within each spot capture region. Enlarged regions encapsulate the outer proliferating ring of the tumor and the immune cell infiltrated core. **B.** Overall percentages of different cell types identified by each deconvolution approach (across all spots combined).

7. **Supplemental Figure S4**

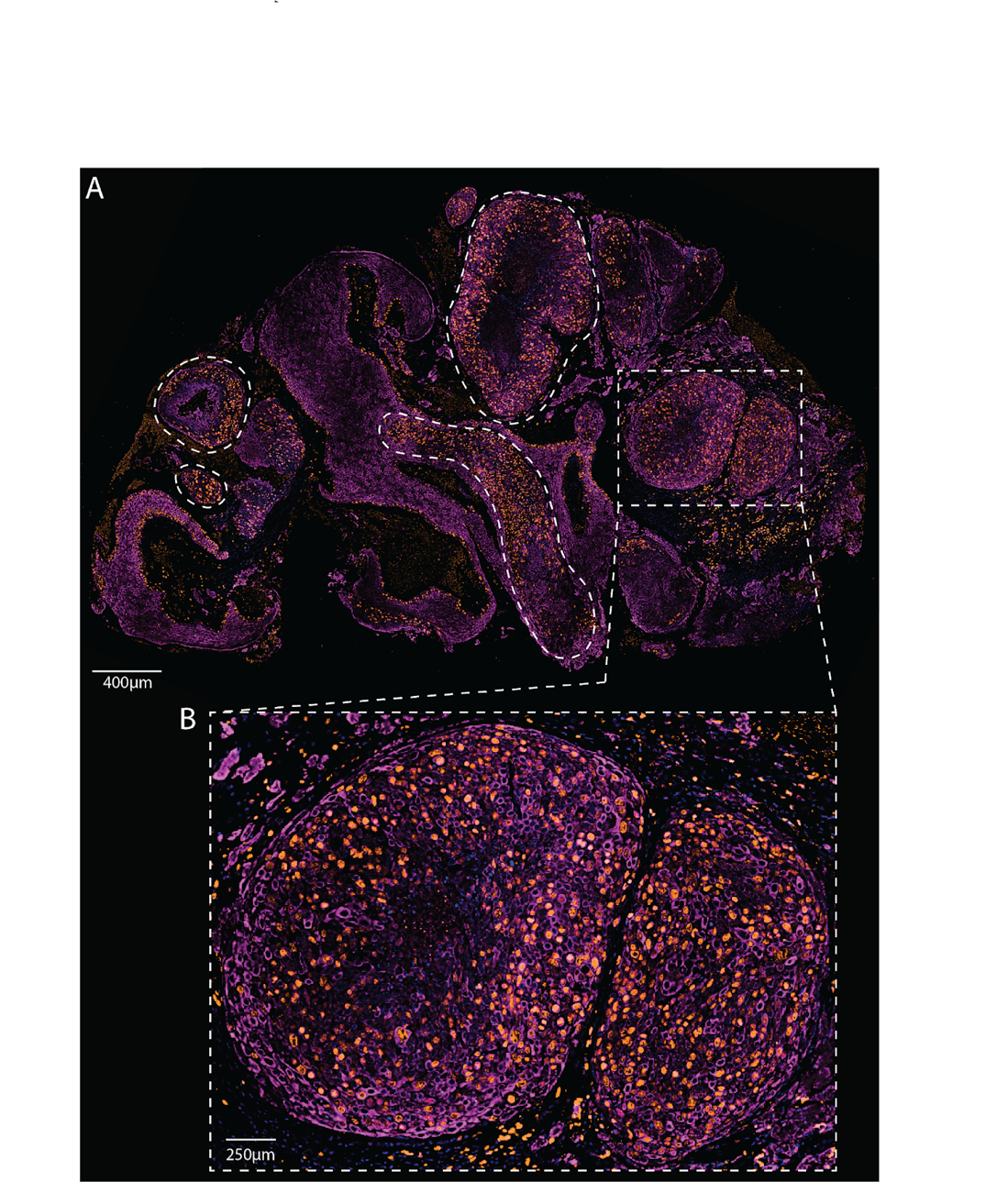

Figure S4. **Localization of proliferating tumor cells. A.** PHENOCYCLER fluorescence images displaying PanCK+ (Purple) and Ki67+ (Orange) cells across the MAR21 OPSCC sample. **B.** Tumor region of interest presents with leading edge of active proliferating cells and inactive tumor core.

8. **Supplemental Figure S5**

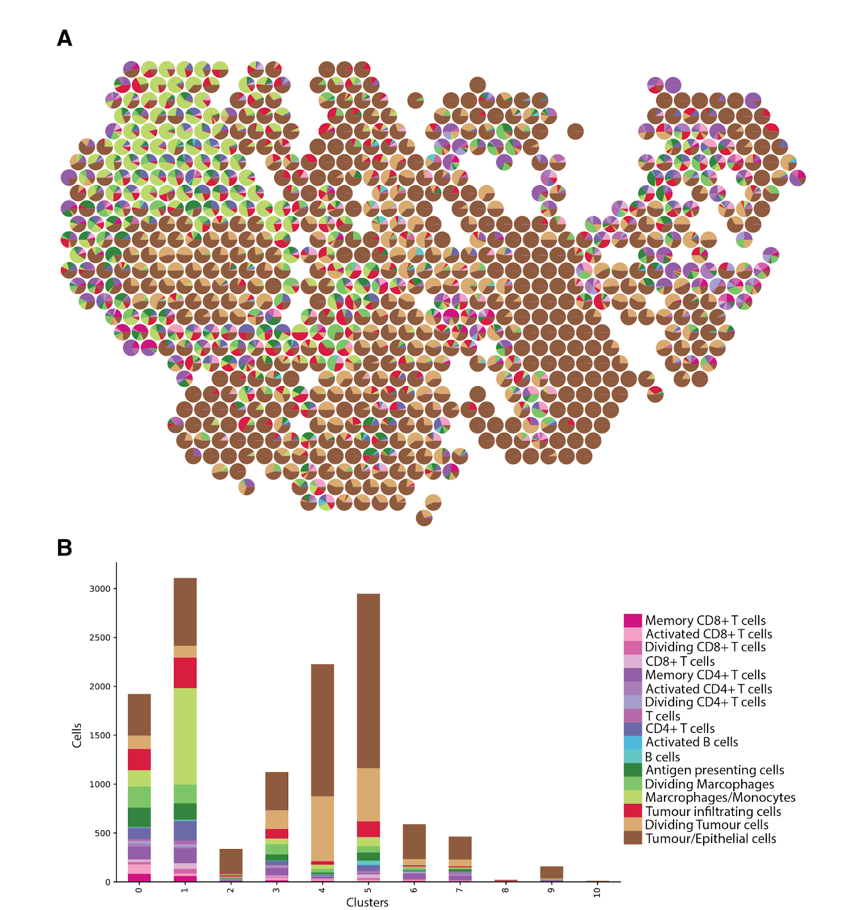

Figure S5. **Integrated Spatial-Omic characterization of tumor immune cell microenvironments. A.** Visium spot deconvolution of MAR21 OPSCC using PHENOCYCLER single-cell data. Spots are presented as pie-charts representing the relative proportions of each cell-type located within each spot. **B.** Quantification of cell-types found within each Visium cluster.

9. **Supplemental Figure S6**

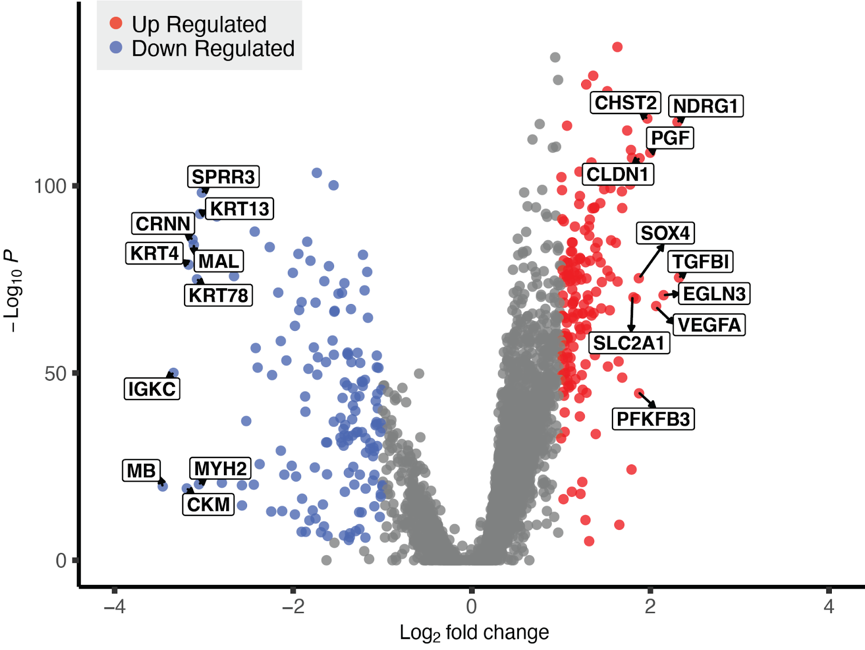

Figure S6. **Transcriptional profile of tumor clusters within MAR21 OPSCC.** DEGs of the combined tumor clusters relative to all other clusters. Significantly up-regulated and down-regulated genes are highlighted in red and blue respectively (p-value < 0.001 and an absolute fold- change > 1). The top 10 over- and under-expressed genes are annotated.

10. **Supplemental Figure S7**

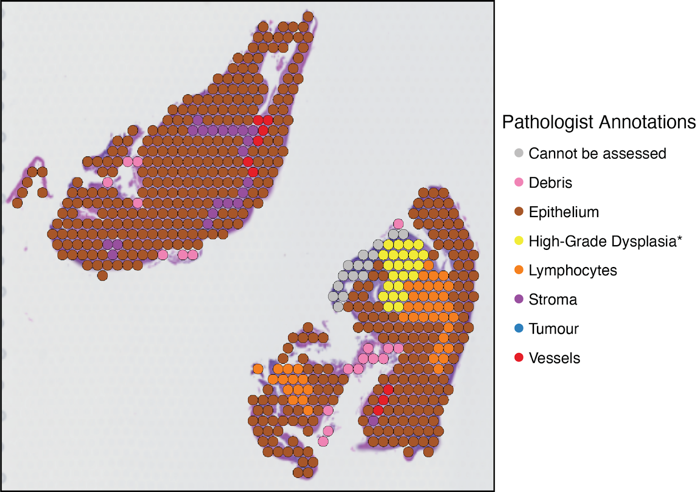

Figure S7. **Initial Pathologist annotation of SEP21 OPSCC sample.** Colored spots identify different morphological tissue structure identified from the H&E image by a clinical pathologist.

11. **Supplemental Figure S8**

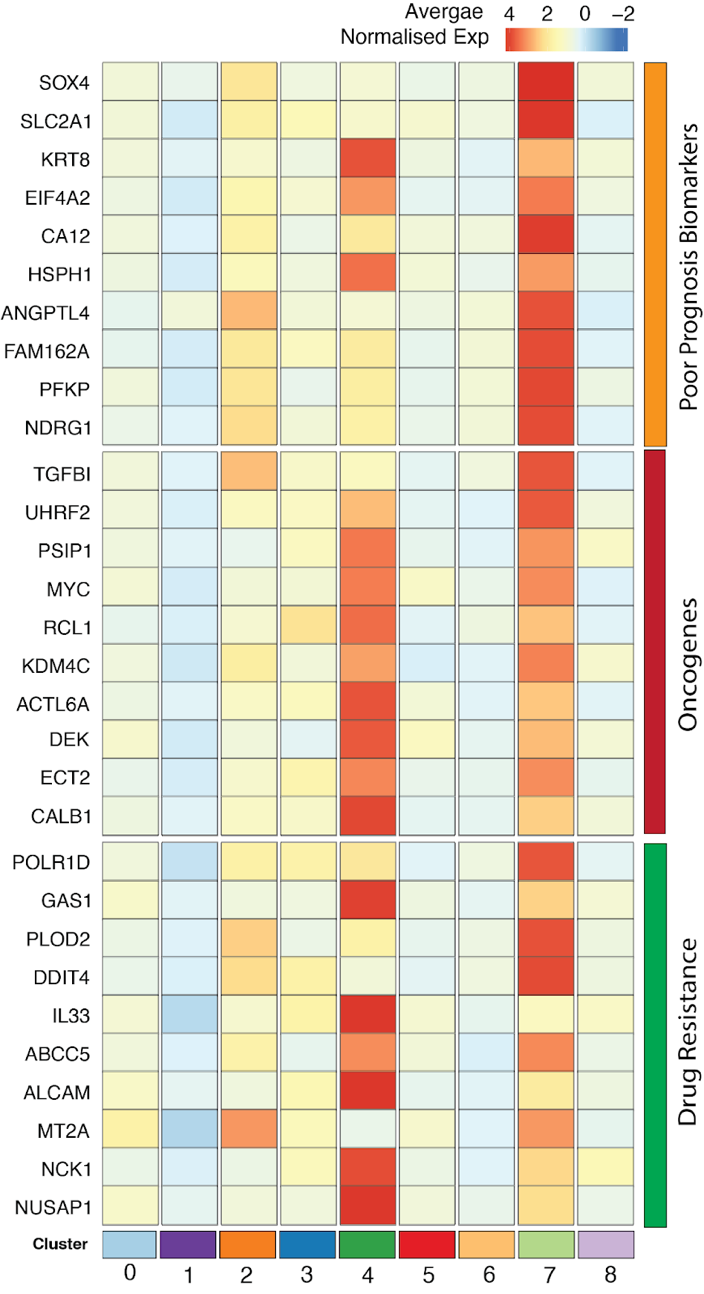

Figure S8. **Tumor transcriptional profile recapitulated in recurrent SEP21 OPSCC.** Relative expression of poor prognosis markers (red), oncogenes (orange) and drug resistance genes (green), previously identified from the combined tumor cluster within MAR21, across each newly defined cluster generated through comparing OPSCC samples. The top 9 genes of each category were displayed. Colors gradient represents average normalized expression values across all spots in each cluster, which were z-transformed by genes (rows of the heatmap).

12. **Supplemental Figure S9**

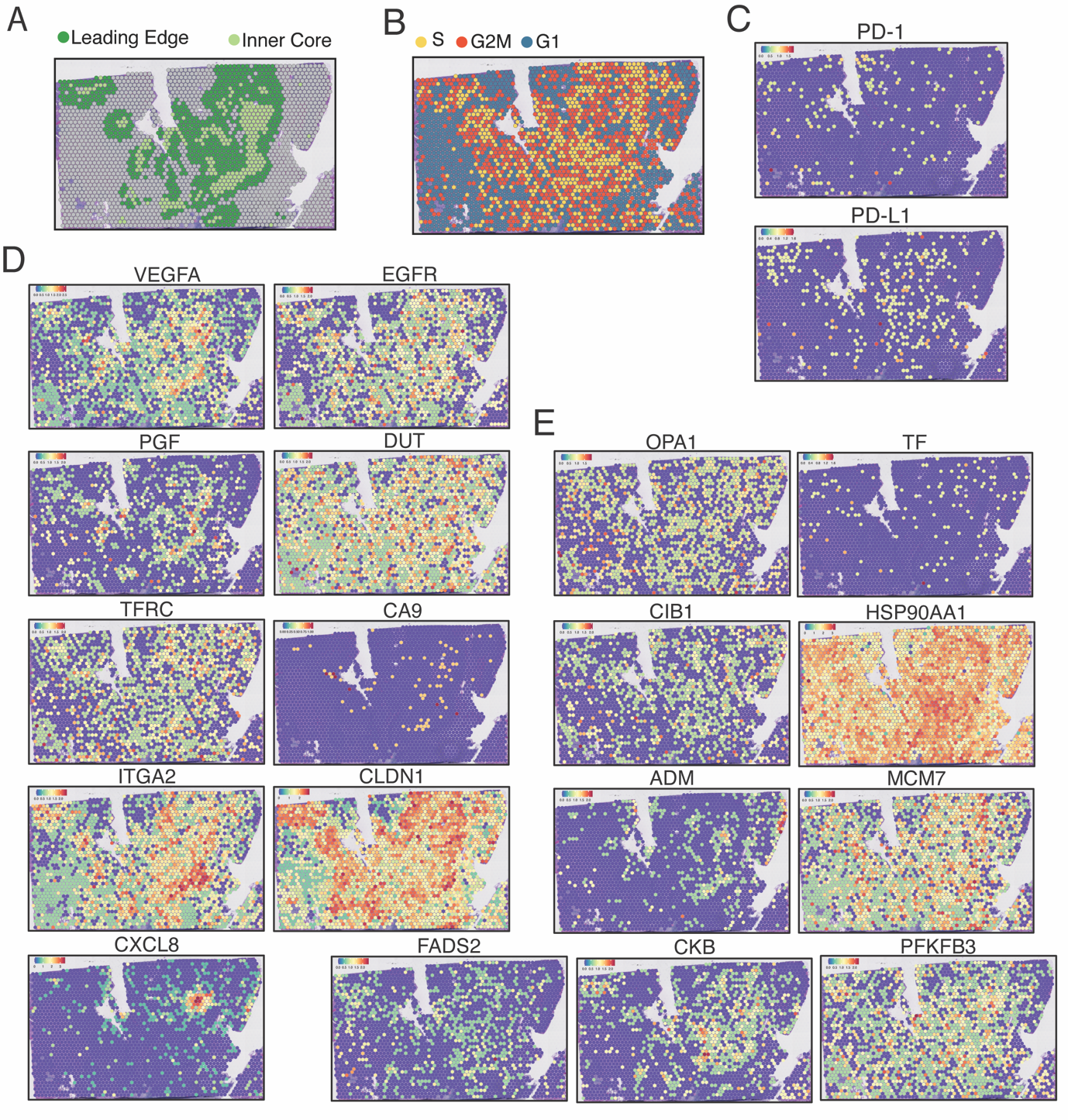

Figure S9. **Top druggable targets differ between additional OPSCC patients.** **A.** Spatial representation of Visium cluster identified as tumor leading edge (dark green) and inner core (light green) within an additional OPSCC patient. **B.** Cell cycle phase of each spot based on relative expression of specific cell phase genes. **C.** Spatial expression of genes targeted by Nivolumab and Pembrolizumab targeted *PD-1/PD-L1* pathway. **D.** Spatial visualization of gene expression of genes targeted by previously identified clinical therapies. **E.** Spatial localization of gene expression levels for select experimental drug therapies.

Table S3. **Gene classification based on function reported in the literature in the cancer setting.**

| **Gene** | **Categories** | **Ref** |
| --- | --- | --- |
| HAUS6 | Oncogenes | ^1^ |
| DBN1 | Oncogenes | ^2-4^ |
| CCNL1 | Oncogenes | ^5^ |
| ACTL6A | Oncogenes | ^6^ |
| CHAF1A | Oncogenes | ^7,8^ |
| DEK | Oncogenes | ^9^ |
| ECT2 | Oncogenes | ^10,11^ |
| EIF4EBP1 | Oncogenes | ^12^ |
| FXR1 | Oncogenes | ^13^ |
| JUN | Oncogenes | ^14^ |
| KDM4C | Oncogenes | ^15^ |
| MYBL2 | Oncogenes | ^16^ |
| MYC | Oncogenes | ^17^ |
| NSD2 | Oncogenes | ^18^ |
| PFDN2 | Oncogenes | ^19^ |
| PSIP1 | Oncogenes | ^20^ |
| RNF19A | Oncogenes | ^21^ |
| TAGLN2 | Oncogenes | ^22,23^ |
| TGFBI | Oncogenes | ^24^ |
| UHRF2 | Oncogenes | ^25^ |
| VAV3 | Oncogenes | ^26,27^ |
| JAG1 | Oncogenes | ^28,29^ |
| CENPW | Oncogenes | ^30,31^ |
| SMC4 | Good and Poor prognosis biomarker | ^32,33^ |
| COL7A1 | Good prognosis biomarker | ^34^ |
| CXADR | Good prognosis biomarker | ^35^ |
| FOXRED2 | Good prognosis biomarker | ^36^ |
| LPAR3 | Good prognosis biomarker | ^37^ |
| MEI1 | Good prognosis biomarker | ^38^ |
| NEFH | Good prognosis biomarker | ^39^ |
| VLDLR | Good prognosis biomarker | ^40^ |
| PGK1 | Poor prognosis biomarker | ^41^ |
| ANGPTL4 | Poor prognosis biomarker | ^42^ |
| ANP32E | Poor prognosis biomarker | ^43^ |
| ATP13A3 | Poor prognosis biomarker | ^44^ |
| BHLHE40 | Poor prognosis biomarker | ^45,46^ |
| CA12 | Poor prognosis biomarker | ^47^ |
| CALB1 | Poor prognosis biomarker | ^48^ |
| CCT3 | Poor prognosis biomarker | ^49-51^ |
| CENPJ | Poor prognosis biomarker | ^52^ |
| CPA4 | Poor prognosis biomarker | ^53^ |
| CSE1L | Poor prognosis biomarker | ^54^ |
| DSC3 | Poor prognosis biomarker | ^55^ |
| DVL3 | Poor prognosis biomarker | ^56^ |
| EIF4A2 | Poor prognosis biomarker | ^57^ |
| FAM162A | Poor prognosis biomarker | ^58^ |
| FAT2 | Poor prognosis biomarker | ^59^ |
| GTF3A | Poor prognosis biomarker | ^60^ |
| HILPDA | Poor prognosis biomarker | ^61^ |
| HSPA1A | Poor prognosis biomarker | ^62^ |
| HSPA1B | Poor prognosis biomarker | ^62^ |
| HSPE1 | Poor prognosis biomarker | ^63^ |
| HSPH1 | Poor prognosis biomarker | ^64^ |
| IER3 | Poor prognosis biomarker | ^65^ |
| KRT8 | Poor prognosis biomarker | ^66^ |
| LAMA5 | Poor prognosis biomarker | ^67,68^ |
| LPCAT1 | Poor prognosis biomarker | ^69,70^ |
| NFIL3 | Poor prognosis biomarker | ^71^ |
| NRARP | Poor prognosis biomarker | ^72,73^ |
| NUP58 | Poor prognosis biomarker | ^74^ |
| NXPH4 | Poor prognosis biomarker | ^75^ |
| PCLAF | Poor prognosis biomarker | ^76^ |
| PFKP | Poor prognosis biomarker | ^77^ |
| PLAUR | Poor prognosis biomarker | ^78^ |
| PLPP2 | Poor prognosis biomarker | ^79^ |
| PNCK | Poor prognosis biomarker | ^80^ |
| PODXL2 | Poor prognosis biomarker | ^81^ |
| PSMD2 | Poor prognosis biomarker | ^82^ |
| PTHLH | Poor prognosis biomarker | ^83^ |
| PUM3 | Poor prognosis biomarker | ^84^ |
| RFC3 | Poor prognosis biomarker | ^85^ |
| RFC4 | Poor prognosis biomarker | ^86^ |
| RHOV | Poor prognosis biomarker | ^21^ |
| RUVBL1 | Poor prognosis biomarker | ^87^ |
| SAT1 | Poor prognosis biomarker | ^88^ |
| SLC2A1 | Poor prognosis biomarker | ^89^ |
| SNAI2 | Poor prognosis biomarker | ^90^ |
| SOD2 | Poor prognosis biomarker | ^91,92^ |
| SOX4 | Poor prognosis biomarker | ^93^ |
| STC2 | Poor prognosis biomarker | ^94,95^ |
| TBL1XR1 | Poor prognosis biomarker | ^96^ |
| TCN1 | Poor prognosis biomarker | ^97^ |
| TCP1 | Poor prognosis biomarker | ^98^ |
| TFDP1 | Poor prognosis biomarker | ^99^ |
| TMEM189 | Poor prognosis biomarker | ^100,101^ |
| U2SURP | Poor prognosis biomarker | ^102^ |
| UBE2C | Poor prognosis biomarker | ^26^ |
| YEATS2 | Poor prognosis biomarker | ^103^ |
| ABCC5 | Drug Resistance | ^104^ |
| ALCAM | Drug Resistance | ^105^ |
| CALCRL | Drug Resistance | ^106,107^ |
| FBXO42 | Drug Resistance | ^108^ |
| GAS1 | Drug Resistance | ^109^ |
| IL33 | Drug Resistance | ^110^ |
| MKNK2 | Drug Resistance | ^111^ |
| MT2A | Drug Resistance | ^112,113^ |
| NCK1 | Drug Resistance | ^114^ |
| NUSAP1 | Drug Resistance | ^115^ |
| PGF | Drug Resistance | ^116^ |
| PLOD2 | Drug Resistance | ^117^ |
| POLR1D | Drug Resistance | ^118^ |
| PGK1 | Drug Resistance | ^41^ |
| NES | Drug Resistance | ^119^ |
| CERS2 | Tumor suppressor | ^120^ |
| DLG1 | Tumor suppressor | ^121^ |
| FLNB | Tumor suppressor | ^122^ |
| GADD45A | Tumor suppressor | ^123^ |
| HLTF | Tumor suppressor | ^124^ |
| HMG20B | Tumor suppressor | ^125^ |
| IL12RB2 | Tumor suppressor | ^126,127^ |
| ILF3 | Tumor suppressor | ^128^ |
| IRF6 | Tumor suppressor | ^129,130^ |
| MT1G | Tumor suppressor | ^131^ |
| MT1X | Tumor suppressor | ^132^ |
| NDRG1 | Tumor suppressor | ^133^ |
| RBM38 | Tumor suppressor | ^134^ |
| RCL1 | Tumor suppressor | ^135^ |
| RSRC1 | Tumor suppressor | ^136^ |
| SEMA4B | Tumor suppressor | ^137^ |
| DDIT4 | Tumor suppressor | ^138,139^ |
| EFNA1 | Tumor suppressor | ^140^ |
| NRG1 | Tumor suppressor | ^141^ |
| ALDH1A3 | Drug Resistance (chemoresistance in colorectal cancer); Experimental target | ^142,143^ |
| HMGB1 | Experimental target | ^144^ |
| CNFN | Good prognosis marker | ^145^ |
| CYSRT1 | Good prognosis marker | ^146,147^ |
| IGKC | Good prognosis marker | ^148^ |
| IGKV4-1 | Good prognosis marker | ^149^ |
| IGLC1 | Good prognosis marker | ^150^ |
| JCHAIN | Good prognosis marker | ^151^ |
| IGHA1 | Good prognosis marker | ^152^ |
| IGHG1 | Good prognosis marker | ^152^ |
| IGHG2 | Good prognosis marker | ^152^ |
| IGHG4 | Good prognosis marker | ^152^ |
| CALB1 | Oncogene | ^48,153^ |
| LCN2 | Oncogene | ^154^ |
| TGFBI | Oncogene | ^24^ |
| XBP1 | Oncogene | ^155^ |
| COL1A1 | Poor prognosis marker | ^156^ |
| COL1A2 | Poor prognosis marker | ^157^ |
| COL3A1 | Poor prognosis marker | ^158^ |
| CRNN | Poor prognosis marker | ^159^ |
| ECM1 | Poor prognosis marker | ^160,161^ |
| EMP1 | Poor prognosis marker | ^162^ |
| ERO1A | Poor prognosis marker | ^163^ |
| GLUL | Poor prognosis marker | ^164,165^ |
| KLK13 | Poor prognosis marker | ^166^ |
| KLK6 | Poor prognosis marker | ^167,168^ |
| KLK7 | Poor prognosis marker | ^168^ |
| KRT13 | Poor prognosis marker | ^169^ |
| KRT4 | Poor prognosis marker | ^170^ |
| KRT78 | Poor prognosis marker | ^171^ |
| KRT8 | Poor prognosis marker | ^171^ |
| LCE3D | Poor prognosis marker | ^172,173^ |
| NDRG1 | Poor prognosis marker | ^174,175^ |
| PRSS27 | Poor prognosis marker | ^176^ |
| SOD2 | Poor prognosis marker | ^91,92^ |
| SPRR2A | Poor prognosis marker | ^177,178^ |
| SPRR2D | Poor prognosis marker | ^178^ |
| SPRR3 | Poor prognosis marker | ^179^ |
| HOPX | Tumor suppressor | ^180,181^ |
| MAL | Tumor suppressor | ^182^ |
| SLURP1 | Tumor suppressor | ^183,184^ |
| SLURP2 | Tumor suppressor | ^183^ |
| SPINK5 | Tumor suppressor | ^185^ |
| SPINK7 | Tumor suppressor | ^186^ |
| TGM3 | Tumor suppressor | ^187,188^ |
| TM4SF1 | Tumor suppressor (gastric carcinoma); poor prognosis marker (epithelial cancers) | ^189,190^ |

### Table S4. **Identified clinical and preclinical targets.**

| **Gene** | **Drug name** | **Company name** | **Indication** | **Development Stage** |
| --- | --- | --- | --- | --- |
| CA9 ^47,191^ | SLC-0111 | SignalChem Lifesciences Corp | Pancreatic Ductal Adenocarcinoma | Phase II |
| DUT ^192^ | TAS-114 | Taiho Pharmaceutical Co Ltd | Adenocarcinoma Of The Gastroesophageal Junction; Gastric Cancer; Non-Small Cell Lung Cancer | Phase II |
|  |  |  | Metastatic Breast Cancer; Metastatic Colorectal Cancer; Pancreatic Cancer | Phase I |
| EGFR ^193^ | futuximab + modotuximab | Symphogen A/S | Metastatic Colorectal Cancer | Phase III |
|  | abivertinib maleate | Sorrento Therapeutics Inc | Metastatic Hormone Refractory (Castration Resistant, Androgen-Independent) Prostate Cancer; Burkitt Lymphoma; Diffuse Large B-Cell Lymphoma; Follicular Lymphoma; Mantle Cell Lymphoma; Marginal Zone B-cell Lymphoma; Refractory Chronic Lymphocytic Leukemia (CLL); Relapsed Chronic Lymphocytic Leukemia (CLL); Waldenstrom Macroglobulinemia (Lymphoplasmacytic Lymphoma) | Phase I |
|  | abivertinib maleate | Sorrento Therapeutics Inc | Hairy Cell Leukemia; Non-Small Cell Lung Cancer; Prostate Cancer | Phase II |
|  | abivertinib maleate | Sorrento Therapeutics Inc | Non-Small Cell Lung Cancer | Phase III |
|  | afatinib dimaleate | More than 10 companies | Non-Small Cell Lung Cancer; Squamous Non-Small Cell Lung Cancer | Marketed |
|  |  | Boehringer Ingelheim International GmbH | Anaplastic Astrocytoma; Anaplastic Oligoastrocytoma; Chordoma; Leptomeningeal Disease (Neoplastic Meningitis, Leptomeningeal Carcinomatosis); Low-Grade Glioma; Medulloblastoma; Meningioma; Oligodendroglioma; Pituitary Tumor; Recurrent Glioblastoma Multiforme (GBM) | Phase I |
|  |  |  | Chordoma; Esophageal Cancer; Gastric Cancer; Metastatic Transitional (Urothelial) Tract Cancer; Squamous Cell Carcinoma; Uterine Cancer; Lymphoma; Refractory Multiple Myeloma | Phase II |
|  | alflutinib mesylate | Allist Shanghai Pharmaceutical Technology Co Ltd | Non-Small Cell Lung Cancer | Marketed |
|  |  |  | Lung Adenocarcinoma | Phase I |
|  |  | Arrivent Biopharma Inc | Non-Small Cell Lung Cancer | Phase II |
|  | amelimumab | Shanghai Sailun Biotechnology Co Ltd | Colorectal Cancer | Phase II |
|  |  | Janssen Inc | Non-Small Cell Lung Cancer | Marketed |
|  |  | Janssen-Cilag Pharma GmbH | Non-Small Cell Lung Cancer | Marketed |
|  |  | Johnson & Johnson | Breast Cancer; Colorectal Cancer; Gastroesophageal (GE) Junction Carcinomas; Head and Neck Cancer Squamous Cell Carcinoma; Hepatocellular Carcinoma; Kidney Cancer (Renal Cell Cancer); Malignant Mesothelioma; Medullary Thyroid Cancer; Non-Small Cell Lung Cancer; Ovarian Cancer; Solid Tumor | Phase I |
|  |  |  | Esophageal Cancer; Gastric Cancer; Gastroesophageal (GE) Junction Carcinomas; Metastatic Colorectal Cancer | Phase II |
|  |  |  | Non-Small Cell Lung Cancer | Phase III |
|  | AMX-3009 | Arromax Pharmatech Co Ltd | Solid Tumor | Phase I |
|  | ASK-120067 | Jiangsu Aosaikang Pharmaceutical Co Ltd | Non-Small Cell Lung Cancer | Phase III |
|  | aumolertinib mesylate | Jiangsu Hansoh Pharmaceutical Group Co Ltd | Non-Small Cell Lung Cancer | Marketed |
|  |  | EQRx Inc | Non-Small Cell Lung Cancer | Phase III |
|  |  | Jiangsu Hansoh Pharmaceutical Group Co Ltd | Non-Small Cell Lung Cancer | Phase III |
|  | BAY-2927088 | Bayer AG | Non-Small Cell Lung Cancer | Phase I |
|  | BBT-176 | Bridge Biotherapeutics Inc | Non-Small Cell Lung Cancer | Phase II |
|  | BC-001 | Dragonboat Biopharmaceutical (Shanghai) Co Ltd | Unspecified Cancer | Phase III |
|  | BCA-101 | Bicara Therapeutics Inc | Anal Cancer; Anaplastic Thyroid Cancer; Colorectal Cancer; Epithelial Ovarian Cancer; Gastric Cancer; Glioblastoma Multiforme (GBM); Head and Neck Cancer Squamous Cell Carcinoma; Hepatocellular Carcinoma; Lung Cancer; Pancreatic Cancer; Solid Tumor; Squamous Cell Carcinoma | Phase II |
|  | BDTX-1535 | Black Diamond Therapeutics Inc | Glioblastoma Multiforme (GBM); Non-Small Cell Lung Cancer | Phase I |
|  | BEBT-109 | Guangzhou BeBetter Medicine Technology Co Ltd | Non-Small Cell Lung Cancer | Phase II |
|  | befortinib mesylate | InventisBio Co Ltd | Non-Small Cell Lung Cancer | Phase III |
|  | BLU-451 | Blueprint Medicines Corp | Non-Small Cell Lung Cancer | Phase II |
|  | BLU-701 | Zai Lab Ltd | Non-Small Cell Lung Cancer | Phase II |
|  | BLU-945 | Blueprint Medicines Corp | Non-Small Cell Lung Cancer | Phase II |
|  | BPI-361175 | Betta Pharmaceuticals Co Ltd | Non-Small Cell Lung Cancer; Solid Tumor | Phase II |
|  | BPI-7711 | Beta Pharma Inc | Non-Small Cell Lung Cancer | Phase II |
|  | C-005 | Wuxi Shuangliang Biotechnology Co Ltd | Non-Small Cell Lung Cancer | Phase I |
|  | cetuximab | Eli Lilly Canada Inc; Merck; Bristol-Myers Squibb KK; PT. Merck Tbk; | Head And Neck Cancer Squamous Cell Carcinoma; Metastatic Colorectal Cancer | Marketed |
|  |  | Merck | Metastatic Colorectal Cancer | Phase I |
|  |  | TheraOp gGmbH | Metastatic Colorectal Cancer | Phase II |
|  |  | Eli Lilly and Co | Anal Cancer | Phase II |
|  |  | Merck | Recurrent Head and Neck Cancer Squamous Cell Carcinoma; Squamous Non-Small Cell Lung Cancer | Phase II |
|  | cetuximab + cobimetinib + palbociclib | Cothera Bioscience Pty Ltd | Metastatic Colorectal Cancer | Phase II |
|  | cetuximab biobetter | Mabpharm Ltd | Metastatic Colorectal Cancer | Phase III |
|  |  | Dragonboat Biopharmaceutical (Shanghai) Co Ltd; Shanghai Jing Ze Biotechnology Co Ltd | Cervical Cancer; Colorectal Cancer; Endometrial Cancer; Esophageal Squamous Cell Carcinoma (ESCC); Head and Neck Cancer Squamous Cell Carcinoma; Metastatic Colorectal Cancer; Ovarian Cancer; Penile Cancer | Phase I |
|  |  | Enzene Biosciences Ltd | Head And Neck Cancer Squamous Cell Carcinoma; Lip Cancer; Locally Recurrent or Locoregional Solid Malignancies; Oral Cavity (Mouth) Cancer; Pharyngeal Neoplasm | Phase III |
|  |  | Ampo Biotechnology Inc; Cinnagen Co; Sichuan Kelun Pharmaceutical Co Ltd; 3SBio Inc | Metastatic Colorectal Cancer | Phase III |
|  |  | R-Pharm | Recurrent Head and Neck Cancer Squamous Cell Carcinoma | Phase III |
|  | dabrafenib mesylate + panitumumab + trametinib dimethyl sulfoxide | Novartis AG | Metastatic Colorectal Cancer | Phase II |
|  | dacomitinib | Pfizer | Non-Small Cell Lung Cancer | Marketed |
|  |  |  | Human Epidermal Growth Factor Receptor 2 Positive Breast Cancer (HER2+ Breast Cancer); Ovarian Cancer; Triple-Negative Breast Cancer (TNBC) | Phase I |
|  | DBPR-112 | AnBogen Therapeutics | Non-Small Cell Lung Cancer; Solid Tumor | Phase II |
|  | DF-203 | Suzhou Dingfu Target Biotechnology Co Ltd | Colorectal Cancer; Head and Neck Cancer Squamous Cell Carcinoma; Non-Small Cell Lung Cancer; Pancreatic Cancer; Renal Cell Carcinoma; Solid Tumor; Triple-Negative Breast Cancer (TNBC) | Phase I |
|  | doxitinib mesylate | Henan Genuine Biotech Co Ltd | Non-Small Cell Lung Cancer | Phase II |
|  | DZD-9008 | Dizal (Jiangsu) Pharmaceutical Co Ltd | Solid Tumor; Non-Small Cell Lung Cancer | Phase II |
|  | EO-1001 | Senz Oncology Pty Ltd | Unspecified Cancer | Phase II |
|  |  | Edison Oncology Holding Corp | Breast Cancer; Central Nervous System (CNS) Cancer; Non-Small Cell Lung Cancer | Phase II |
|  | epertinib | Shionogi & Co Ltd | Breast Cancer; Malignant Neoplasms | Phase II |
|  |  | EOC Pharma Ltd | Breast Cancer | Phase II |
|  |  | Hutchison MediPharma Ltd | Glioblastoma Multiforme (GBM) | Phase II |
|  | ERAS-801 | Erasca Inc | Recurrent Glioblastoma Multiforme (GBM) | Phase I |
|  | erlotinib | More than 15 companies | Metastatic Pancreatic Cancer; Non-Small Cell Lung Cancer | Marketed |
|  | erlotinib hydrochloride | More than 50 companies | Metastatic Pancreatic Cancer; Non-Small Cell Lung Cancer | Marketed |
|  | ES-072 | Zhejiang Bossan Pharmaceutical Co Ltd | Non-Small Cell Lung Cancer | Phase I |
|  | FCN-411 | Fochon Pharmaceutical Ltd | Head And Neck Cancer Squamous Cell Carcinoma | Phase I |
|  |  |  | Non-Small Cell Lung Cancer | Phase II |
|  | FHND-9041 | Jiangsu Zhengda Fenghai Pharmaceutical Co Ltd | Non-Small Cell Lung Cancer | Phase III |
|  | Fusion Protein to Antagonize EGFR for Glioblastoma Multiforme and Malignant Glioma | Istari Oncology Inc | Glioblastoma Multiforme (GBM); Malignant Glioma | Phase II |
|  | FWD-1509 | Shenzhen Forward Pharmaceutical Co Ltd | Non-Small Cell Lung Cancer | Phase II |
|  | GB-263 | Genor BioPharma Co Ltd | Non-Small Cell Lung Cancer; Solid Tumor; Esophageal Cancer; Gastric Cancer; Head and Neck Cancer Squamous Cell Carcinoma; Metastatic Colorectal Cancer | Phase II |
|  | GC-1118A | GC Biopharma Corp | Adenocarcinoma Of the Gastroesophageal Junction; Colon Cancer; Gastric Cancer; Metastatic Colorectal Cancer; Solid Tumor | Phase II |
|  | gefitinib | AstraZeneca; Daiichi Sankyo Espha Co Ltd | Non-Small Cell Lung Cancer | Marketed |
|  | H-002 | RedCloud Bio Inc | Non-Small Cell Lung Cancer | Phase II |
|  | HL-07 | Hualan Biological Engineering Inc | Metastatic Colorectal Cancer | Phase II |
|  | HS-627 | Zhejiang Hisun Pharmaceutical Co Ltd | Triple-Negative Breast Cancer (TNBC) | Phase I |
|  |  |  | Human Epidermal Growth Factor Receptor 2 Positive Breast Cancer (HER2+ Breast Cancer) | Phase III |
|  | icotinib hydrochloride | Betta Pharmaceuticals Co Ltd | Non-Small Cell Lung Cancer | Marketed |
|  | JMT-101 | CSPC Pharmaceutical Group Ltd | Esophageal Squamous Cell Carcinoma (ESCC); Metastatic Colorectal Cancer; Non-Small Cell Lung Cancer; Solid Tumor | Phase I |
|  |  |  | Non-Small Cell Lung Cancer | Phase II |
|  | JS-111 | Shanghai Junshi Bioscience Co Ltd | Non-Small Cell Lung Cancer | Phase II |
|  | KBP-5209 | XuanZhu Biological Technology Co Ltd | Breast Cancer; Carcinoma of Unknown Primary (Occult Primary Tumor/Cancer of Unknown Primary); Colorectal Cancer; Gallbladder Cancer; Gastric Cancer; Head and Neck Cancer; Non-Small Cell Lung Cancer; Ovarian Cancer; Pancreatic Cancer; Paranasal Sinus And Nasal Cavity Cancer; Sarcomas | Phase II |
|  | lapatinib | Lupin Pharmaceuticals Inc; Sayre Therapeutics | Human Epidermal Growth Factor Receptor 2 Positive Breast Cancer (HER2+ Breast Cancer) | Marketed |
|  | lapatinib ditosylate | Zhuhai Rundu Pharmaceutical Co Ltd; Hetero Healthcare Ltd; Cipla Ltd | Human Epidermal Growth Factor Receptor 2 Positive Breast Cancer (HER2+ Breast Cancer) | Marketed |
|  | larotinib | HEC Pharma Co Ltd | Esophageal Squamous Cell Carcinoma (ESCC) | Phase III |
|  |  | Yuhan Corp | Non-Small Cell Lung Cancer | Marketed |
|  |  | Genosco Inc | Metastatic Brain Tumor | Phase II |
|  |  | Johnson & Johnson; Genosco Inc | Non-Small Cell Lung Cancer | Phase III |
|  | lifirafenib maleate | BeiGene Ltd | Colorectal Cancer; Endometrial Cancer; Non-Small Cell Lung Cancer; Ovarian Cancer; Pancreatic Cancer; Cholangiocarcinoma; Melanoma; Papillary Thyroid Cancer; | Phase II |
|  | LL-191 | Nalo Therapeutics Inc | Non-Small Cell Lung Cancer | Phase I |
|  | MCLA-129 | Merus NV; Betta Pharmaceuticals Co Ltd | Adenocarcinoma Of the Gastroesophageal Junction; Esophageal Squamous Cell Carcinoma (ESCC); Gastric Cancer; Head And Neck Cancer Squamous Cell Carcinoma; Non-Small Cell Lung Cancer | Phase II |
|  | MET-306 | Huadong Medicine Co Ltd | Non-Small Cell Lung Cancer | Phase III |
|  | mobocertinib | Takeda Pharmaceuticals Australia Pty Ltd | Non-Small Cell Lung Cancer | Marketed |
|  |  |  | Bladder Cancer; Breast Cancer; Esophageal Cancer; Gastric Cancer; Head and Neck Cancer; Metastatic Biliary Tract Cancer; Urinary Tract Cancer | Phase II |
|  |  |  | Non-Small Cell Lung Cancer | Phase III |
|  | MVC-101 | Maverick Therapeutics Inc | Colorectal Cancer; Head and Neck Cancer Squamous Cell Carcinoma; Non-Small Cell Lung Cancer; Pancreatic Cancer; Solid Tumor | Phase II |
|  | naquotinib mesylate | Novartis AG | Non-Small Cell Lung Cancer | Phase II |
|  | necitumumab | Nippon Kayaku Co Ltd; Eli Lilly Canada Inc  Biotech Pharmaceutical Co Ltd  PT Kalbe Farma Tbk | Squamous Non-Small Cell Lung Cancer | Marketed |
|  |  |  | Nasopharyngeal Cancer | Marketed |
|  |  |  | High-Grade Glioma | Marketed |
|  |  | Biocon Ltd | Head And Neck Cancer Squamous Cell Carcinoma | Marketed |
|  |  | Eurofarma Laboratorios SA | Glioma; High-Grade Glioma; Pediatric Diffuse Intrinsic Pontine Glioma | Marketed |
|  |  | Biotech Pharmaceutical Co Ltd | Recurrent Head and Neck Cancer Squamous Cell Carcinoma | Phase I |
|  |  | Biotech Pharmaceutical Co Ltd | Head And Neck Cancer Squamous Cell Carcinoma | Phase II |
|  |  | InnoMab Pte Ltd | Cervical Cancer | Phase III |
|  |  | Biotech Pharmaceutical Co Ltd | Pediatric Diffuse Intrinsic Pontine Glioma | Phase III |
|  | NRC-2694 | Natco Pharma Ltd | Hypopharyngeal Cancer; Laryngeal Cancer; Oral Cavity (Mouth) Cancer; Oropharyngeal Cancer; Recurrent Head and Neck Cancer Squamous Cell Carcinoma | Phase II |
|  | olafertinib | Checkpoint Therapeutics Inc; Suzhou Neupharma Co Ltd | Non-Small Cell Lung Cancer | Phase II |
|  | olmutinib hydrochloride | Hanmi Pharmaceuticals Co Ltd | Non-Small Cell Lung Cancer | Marketed |
|  | osimertinib mesylate | AstraZeneca | Non-Small Cell Lung Cancer | Marketed |
|  | panitumumab | Amgen; PT Glaxo Wellcome; Takeda Pharmaceutical Co Ltd; Amgen Inc; Dr. Reddy's Laboratories Ltd | Metastatic Colorectal Cancer | Marketed |
|  | PB-357 | Puma Biotechnology Inc | Breast Cancer | Phase I |
|  | petosemtamab | Merus NV | Metastatic Colorectal Cancer; Solid Tumor | Phase II |
|  | poziotinib hydrochloride | Hanmi Pharmaceuticals Co Ltd | Esophageal Cancer | Phase I |
|  |  |  | Human Epidermal Growth Factor Receptor 2 Positive Breast Cancer (HER2+ Breast Cancer); Lung Adenocarcinoma | Phase II |
|  |  |  | Non-Small Cell Lung Cancer | Phase III |
|  | pyrotinib | Jiangsu Hengrui Medicine Co Ltd | Human Epidermal Growth Factor Receptor 2 Positive Breast Cancer (HER2+ Breast Cancer) | Marketed |
|  | QL-1105 | Qilu Pharmaceutical Co Ltd | Colorectal Cancer; Head and Neck Cancer | Phase I |
|  | QL-1203 | Qilu Pharmaceutical Co Ltd | Metastatic Colorectal Cancer | Phase III |
|  | RXDX-105 | F. Hoffmann-La Roche Ltd | Leptomeningeal Disease (Neoplastic Meningitis, Leptomeningeal Carcinomatosis); Lung Adenocarcinoma; Medullary Thyroid Cancer; Ovarian Cancer; Solid Tumor; Squamous Non-Small Cell Lung Cancer | Phase I |
|  | sapitinib | AstraZeneca Plc | Metastatic Colorectal Cancer | Phase III |
|  | SCT-200 | SinoCelltech Group Ltd | Colorectal Cancer | Phase I |
|  |  |  | Recurrent Head and Neck Cancer Squamous Cell Carcinoma | Phase II |
|  |  |  | Esophageal Squamous Cell Carcinoma (ESCC); Metastatic Colorectal Cancer; Triple-Negative Breast Cancer (TNBC) | Phase II |
|  | selatinib ditosilate | Qilu Pharmaceutical Co Ltd | Breast Cancer | Phase I |
|  | sirotinib | XuanZhu Biological Technology Co Ltd | Esophageal Squamous Cell Carcinoma (ESCC); Gastric Cancer; Lung Cancer | Phase I |
|  | SKLB-1028 | CSPC Pharmaceutical Group Ltd | Acute Myelocytic Leukemia (AML, Acute Myeloblastic Leukemia) | Phase II |
|  |  |  | Refractory Acute Myeloid Leukemia; Relapsed Acute Myeloid Leukemia | Phase III |
|  | SPH-118811 | Shanghai Pharmaceutical Group Co Ltd | Non-Small Cell Lung Cancer | Phase I |
|  | Sutetinib maleate | Suzhou Teligene Ltd | Non-Small Cell Lung Cancer | Phase II |
|  | SYHA-12128 | CSPC Pharmaceutical Group Ltd | Adenocarcinoma Of the Gastroesophageal Junction; Colorectal Cancer; Gastric Cancer | Phase I |
|  |  |  | Bile Duct Cancer (Cholangiocarcinoma); Extrahepatic Bile Duct Cancer; Gallbladder Cancer; Metastatic Biliary Tract Cancer; Medullary Thyroid Cancer; Non-Small Cell Lung Cancer | Phase II |
|  | SYN-004 | Synermore Biologics Co Ltd | Head And Neck Cancer Squamous Cell Carcinoma; Metastatic Colorectal Cancer; Non-Small Cell Lung Cancer; Solid Tumor | Phase I |
|  | TAS-2940 | Taiho Oncology Inc | Glioblastoma Multiforme (GBM); Human Epidermal Growth Factor Receptor 2 Positive Breast Cancer (HER2+ Breast Cancer); Non-Small Cell Lung Cancer | Phase I |
|  | tesevatinib tosylate | Kadmon Holdings Inc | Leptomeningeal Disease (Neoplastic Meningitis, Leptomeningeal Carcinomatosis); Metastatic Brain Tumor; Recurrent Glioblastoma Multiforme (GBM); Non-Small Cell Lung Cancer | Phase II |
|  | tomuzotuximab | Glycotope GmbH | Breast Cancer; Esophageal Cancer; Gastric Cancer; Gynecological Cancer; Kidney Cancer (Renal Cell Cancer); Metastatic Colorectal Cancer; Non-Small Cell Lung Cancer | Phase I |
|  |  |  | Recurrent Head and Neck Cancer Squamous Cell Carcinoma | Phase II |
|  | TQB-3804 | Chia Tai Tianqing Pharmaceutical Group Co Ltd | Malignant Neoplasms | Phase I |
|  | Vaccine | Center of Molecular Immunology | Hormone Refractory (Castration Resistant, Androgen-Independent) Prostate Cancer | Phase I |
|  | vandetanib | Sanofi-Aventis (Suisse); Medley Industria Farmaceutica Ltda; Genzyme Europe BV | Medullary Thyroid Cancer | Marketed |
|  | varlitinib | Aslan Pharmaceuticals Ltd | Bladder Cancer; Breast Cancer; Colorectal Cancer; Gastrointestinal Tumor; Glioblastoma Multiforme (GBM); Head and Neck Cancer Squamous Cell Carcinoma; Hepatobiliary System Tumor; Metastatic Hepatocellular Carcinoma (HCC); Non-Small Cell Lung Cancer; Ovarian Cancer; Pancreatic Cancer; Prostate Cancer | Phase I |
|  |  |  | Gastric Cancer; Human Epidermal Growth Factor Receptor 2 Positive Breast Cancer (HER2+ Breast Cancer); Gallbladder Cancer | Phase II |
|  |  |  | Gastroesophageal (GE) Junction Carcinomas; Cholangiocarcinoma | Phase III |
|  |  |  | Papillary Thyroid Cancer; Solid Tumor | Phase III |
|  | VRN-07 | Voronoi Group | Solid Tumor | Phase I |
|  | WJ-13404 | Wigen Biomedicine Technology (Shanghai) Co Ltd | Non-Small Cell Lung Cancer | Phase II |
|  | WSD-0922 | Wayshine Biopharma Inc | Non-Small Cell Lung Cancer; Anaplastic Astrocytoma; Glioblastoma Multiforme (GBM) | Phase I |
|  | XZP-5809 | Sihuan Pharmaceutical Holdings Group Ltd | Non-Small Cell Lung Cancer | Phase I |
|  | yinlitinib | HEC Pharma Co Ltd | Breast Cancer | Phase I |
|  | YZJ-0318 | Yangtze River Pharmaceutical Group | Non-Small Cell Lung Cancer | Phase I |
|  | zipalertinib | Taiho Pharmaceutical Co Ltd; Zai Lab Ltd | Non-Small Cell Lung Cancer | Phase II |
|  | ZNE-4 | Zentalis Pharmaceuticals Inc | Non-Small Cell Lung Cancer | Phase II |
|  | zorifertinib | Alpha Biopharma Ltd | Non-Small Cell Lung Cancer | Phase III |
| PGF ^116^ | conbercept | Chengdu Kanghong Pharmaceuticals Group Co Ltd | Retinoblastoma | Phase II |
|  | ziv-aflibercept | Sanofi-Aventis | Metastatic Colorectal Cancer | Marketed |
|  | ziv-aflibercept | Regeneron Pharmaceuticals Inc | Uveal Melanoma | Phase II |
|  | THR-317 | Oncurious NV | Medulloblastoma | Phase II |
|  | ziv-aflibercept biosimilar | Luye Pharma Group Ltd | Metastatic Colorectal Cancer | Phase II |
| CXCL8 ^194^ | BMS-986253 | Bristol-Myers Squibb Co | Head And Neck Cancer Squamous Cell Carcinoma; Hepatocellular Carcinoma; Hormone-Sensitive Prostate Cancer; Melanoma; Non-Small Cell Lung Cancer; Solid Tumor | Phase II |
|  |  |  | Melanoma; Renal Cell Carcinoma | Phase I |
| ITGA2 ^195^ | E-7820 | Eisai Co Ltd | Chronic Myelomonocytic Leukemia (CMML); Myelodysplastic Syndrome; Refractory Acute Myeloid Leukemia; Relapsed Acute Myeloid Leukemia | Phase II |
| HK2 ^196^ | tuvatexib | Vidac Pharma | Cutaneous T-Cell Lymphoma | Phase II |
| HMGB1 ^197^ | dociparstat sodium | Chimerix Inc | Acute Myelocytic Leukemia (AML, Acute Myeloblastic Leukemia) | Phase III |
| CLDN1 ^198^ | ALEF-02 | Alentis Therapeutics AG | Solid Tumor | Phase I |
| VEGFA ^199^ | ABL-001 | Compass Therapeutics Inc | Extrahepatic Bile Duct Cancer; Gallbladder Cancer; Metastatic Biliary Tract Cancer; Metastatic Colorectal Cancer | Phase I |
|  |  | Handok Inc; Elpiscience Biopharma Ltd | Bile Duct Cancer (Cholangiocarcinoma); Extrahepatic Bile Duct Cancer; Gallbladder Cancer; Gastric Cancer; Gastrointestinal Stromal Tumor (GIST); Metastatic Colorectal Cancer; Non-Small Cell Lung Cancer; Ovarian Cancer; Pancreatic Cancer; Solid Tumor | Phase II |
|  | bevacizumab | Roche; Chugai Pharmaceutical Co Ltd; Genentech; Cipla; PT Boehringer Ingelheim | Cervical Cancer; Epithelial Ovarian Cancer; Fallopian Tube Cancer; Metastatic Breast Cancer; Metastatic Colorectal Cancer; Metastatic Renal Cell Carcinoma; Non-Small Cell Lung Cancer; Peritoneal Cancer | Marketed |
|  | bevacizumab | Chugai Pharmaceutical Co Ltd; F. Hoffmann-La Roche Ltd | Hepatocellular Carcinoma; Human Epidermal Growth Factor Receptor 2 Negative Breast Cancer (HER2- Breast Cancer) | Phase II |
|  | bevacizumab | Chugai Pharmaceutical Co Ltd; F. Hoffmann-La Roche Ltd | Cervical Cancer; Malignant Pleural Mesothelioma; Melanoma; Hepatocellular Carcinoma; Metastatic Colorectal Cancer; Lung Cancer; Human Epidermal Growth Factor Receptor 2 Negative Breast Cancer (HER2- Breast Cancer) | Phase III |
|  | bevacizumab + paclitaxel | Sorrento Therapeutics Inc | Adenocarcinoma; Cervical Cancer; Clear Cell Squamous Cell Carcinoma; Endometrial Cancer; Fallopian Tube Cancer; Melanoma; Ovarian Cancer; Peritoneal Cancer; Squamous Cell Carcinoma | Phase I |
|  | bevacizumab biosimilar | Pfizer; Amgen Inc; Daiichi Sankyo Co Ltd; PT Pyridam Farma Tbk; BeiGene Ltd; Cipla; Celltrion Inc; Eris Lifesciences Ltd; Biocad; Qilu Pharmaceutical Co Ltd; Merck | Epithelial Ovarian Cancer; Fallopian Tube Cancer; Glioblastoma Multiforme (GBM); Metastatic Colorectal Cancer; Non-Small Cell Lung Cancer; Peritoneal Cancer; Cervical Cancer; Fallopian Tube Cancer; Metastatic Breast Cancer; Metastatic Renal Cell Carcinoma; Peritoneal Cancer | Marketed |
|  | ziv-aflibercept | Sanofi-Aventis | Metastatic Colorectal Cancer | Marketed |
|  |  | Regeneron Pharmaceuticals Inc | Uveal Melanoma | Phase II |
|  | ziv-aflibercept biosimilar | Luye Pharma Group Ltd | Metastatic Colorectal Cancer | Phase II |
| TFRC | CX-2029 | CytomX Therapeutics Inc | Diffuse Large B-Cell Lymphoma; Squamous Non-Small Cell Lung Cancer.  Adenoid Cystic Carcinoma (ACC); Bladder Cancer; Colorectal Cancer; Esophageal Cancer; Gastroesophageal (GE) Junction Carcinomas; Head and Neck Cancer Squamous Cell Carcinoma; Hepatocellular Carcinoma; Kidney Cancer (Renal Cell Cancer); Malignant Pleural Mesothelioma; Ocular Melanoma; Ovarian Cancer; Pancreatic Cancer; Prostate Cancer; Soft Tissue Sarcoma; Solid Tumor; Thymic Carcinoma; Thymoma (Thymic Epithelial Tumor); Thyroid Cancer | Phase II |
|  | INA-03 | Inatherys | Acute Lymphocytic Leukemia (ALL, Acute Lymphoblastic Leukemia); Acute Myelocytic Leukemia (AML, Acute Myeloblastic Leukemia); Leukemia; Refractory Acute Myeloid Leukemia; Relapsed Acute Myeloid Leukemia | Phase I |
|  | JSTTFR-09 | Fujifilm Holdings Corp | Polycythemia Vera | Phase I |
| PFKFB3 ^200^ | KAN-0438757 | Kancera AB | Triple-Negative Breast Cancer (TNBC) | Preclinical |
| CIB1 ^201-203^ | Small Molecules | Reveris Therapeutics LLC | Breast Cancer | Preclinical |
| HSP90AA1 ^204^ | Small Molecule | China Medical University | Unspecified Cancer | Preclinical |
